# A Barley Powdery Mildew Fungus Non-Autonomous Retrotransposon Encodes a Peptide that Supports Penetration Success on Barley

**DOI:** 10.1101/242271

**Authors:** Mathias Nottensteiner, Bernd Zechmann, Christopher McCollum, Ralph Hückelhoven

## Abstract

Plant immunity is overcome by pathogens by the means of secreted effectors. Host effector targets might be proteins acting in pathogen defense or serve demands of the pathogen. The barley ROP GTPase HvRACB is involved in entry of the powdery mildew fungus *Blumeria graminis* f.sp. *hordei (Bgh)* into barley epidermal cells. We found that HvRACB interacts with the <u>ROP</u>-<u>i</u>nteractive peptide 1 (ROPIP1) that is encoded on the active non-long terminal repeat retroelement Eg-R1 of *Bgh*. Over-expression of ROPIP1 in barley epidermal cells and host-induced post-transcriptional gene silencing (HIGS) of *ROPIP1* suggested that ROPIP1 is involved in virulence of *Bgh*. Bimolecular fluorescence complementation and co-localization supported that ROPIP1 can interact with activated HvRACB in planta. We show that ROPIP1 is expressed by *Bgh* on barley and translocated into the cytoplasm of infected barley cells. ROPIP1 is recruited to microtubules upon co-expression of MICROTUBULE ASSOCIATED ROP GTPase ACTIVATING PROTEIN (HvMAGAP1) and can destabilize cortical microtubules. *Bgh* ROPIP might target HvRACB and manipulate host cell microtubule organization for facilitated host cell entry. Data suggest a possible neo-functionalization of retroelement-derived transcripts for the evolution of a pathogen virulence effector.

## INTRODUCTION

Lots of effort is invested in the understanding of plant immunity against infection by pathogens (Spoel and Dong, 2012) and the underlying genes like Resistance-genes (*R*-genes) or quantitative trait loci (QTL) that might be used in breeding for crops with improved resistance.

In the general model, plant immunity towards invading pathogens is composed of two-main layers, which are pattern-triggered immunity (Macho and Zipfel, 2014) and effector-triggered immunity (Spoel and Dong, 2012). Adapted pathogens evolved means to overcome host immunity, which is mainly attributed to secreted effector proteins that manipulate host cellular processes to the profit of the pathogen. Plant hosts on the other hand evolved resistance proteins that either directly recognize the presence of a corresponding effector or indirectly recognize the action of effector proteins on their host targets or on host decoy proteins that molecularly mimic host targets. Resistance protein signaling accelerates and increases defense responses typically resulting in in the hypersensitive cell death response thereby restricting further proliferation of biotrophic and hemibiotropic pathogens. The exerted mutual selection pressure drives co-evolution of host *R*-genes and pathogen effectors (Jones and Dangl, 2006).

The investigation of host-factors that allow susceptibility against a pathogen is an alternative approach to searching for factors of host immunity. The products of susceptibility genes (*S*-genes) might function in the regulation of plant defense responses or cell death. Alternatively, susceptibility factors can be part of essential cellular processes from which the pathogen profits or that get co-opted by pathogens. The loss of function of *S*-gene products bears the chance of durable pathogen control due to the loss of a cellular function required for compatibility, given that possible pleiotropic effects are not detrimental for plant cultivation (see (van Schie and Takken, 2014) for review). A paradigm example for making use of the loss of *S*-gene functionality is the *MLO* gene, that represents a negative regulator of basal resistance against powdery mildews. Loss of MLO function is associated with powdery mildew resistance in diverse commercially important crop plant species (Kusch and Panstruga, 2017).

The ascomycete *Blumeria graminis* f.sp.*hordei* (*Bgh*) grows and reproduces on living host tissue where it causes barley powdery mildew. It forms an appressorium and an infection peg for penetration of the host epidermal at 10–15 hours after inoculation (hai). The penetrating hyphae differentiates into a mature haustorium until 48 hai. Haustoria stay separated from the host cell cytoplasm by the extrahaustorial matrix and a surrounding host membrane, the extrahaustorial membrane. Besides expanding the surface for absorption of carbohydrates and amino acids (Voegele et al., 2001), haustoria serve effector delivery into host cells. Penetration of the host cell is a prerequisite for further epicuticular development and asexual reproduction of *Bgh*.

The genomes of *Bgh* and of the close relative *Blumeria graminis f.sp. tritici* (*Bgt*) are sequenced (Spanu et al., 2010; Wicker et al., 2013). Effector proteins of *Blumeria graminis* are either identified via their avirulence (Avr) function if they are recognized by corresponding R proteins or because of canonical characteristics of secreted effector proteins. *Bgh* encodes more than 500 candidate secreted effector proteins (CSEPs) (Pedersen et al., 2012) identified by defined criteria for effector architecture. Some CSEPs are alternatively called BECs for *Blumeria* effector candidates, if they have been found to be expressed in *Bgh*-infected barley tissue (Bindschedler et al., 2009; Pliego et al., 2013). Recently, several CSEP proteins were shown to act as Avr factors in race-specific resistance of wheat and barley (Bourras et al., 2015; Lu et al., 2016; Praz et al., 2017). *Bgh* also encodes 1350 paralogous copies of the second class of *Bgh* effector candidates, EKAs (Effectors homologous to *Avrk1* and *Avra10*), which do not encode N-terminal signal peptides. The EKAs Avr_a10_ and Avr_k1_ are reported to be recognized by the corresponding barley R-proteins MLA10 and MLK1, respectively (Nowara et al., 2010; Ridout et al., 2006; Shen et al., 2007). Avr_a10_ and Avr_k1_ evolved from 3’-truncated ORF1 proteins of *Bgh* long-interspersed element (LINE) retrotransposons (Amselem et al., 2015). The ~120Mb genome of *Bgh* and other powdery mildews showed highly enlarged genome sizes in comparison to the ascomycete mean which was attributed to high transposable element (TE) activity. The genome of *Bgh* was estimated to be composed of ~65 % TEs, respectively ~75 % repetitive DNA content in total (Spanu et al., 2010) and > 90 % repetitive DNA content were estimated for *Bgt* of wheat (Wicker et al., 2013). Both species showed a substantial loss in gene number including genes for enzymes of the primary and secondary metabolism. This might reflect their adaption to their obligate biotrophic lifestyle and a minimum key gene set, of which deletion would be detrimental for fitness.

The bulk of TE-content in the *Bgh* genome are class I retrotransposons. Thereof non-long-terminal-repeat (LTR) retrotransposons are more abundant than the retrovirus-related LTR-retrotransposons. Within Non-LTR retrotransposons, autonomous LINE elements are more abundant than non-autonomous short-interspersed elements (SINEs) that typically need LINE assistance for retro-transposition as they do not encode the required proteins. The SINE-classified Non-LTRs Eg-R1 (Wei et al., 1996) and Egh24 (Rasmussen et al., 1993) e.g. cover ca. 10 % of the *Bgh* genome space (Spanu et al., 2010).

The *Hordeum vulgare* (*Hv*) small monomeric Rho of plants (ROP) GTPase HvRACB has been shown to support *Bgh* haustorial ingrowth into barley epidermal cells when expressed as constitutively activated (CA) mutant (Scheler et al., 2016; Schultheiss et al., 2003). *Vice versa* RNA interference (RNAi) mediated silencing of *HvRACB* in barley restricts haustorial invasion (Hoefle et al., 2011; Scheler et al., 2016; Schultheiss et al., 2002). The abundance of activated GTP-bound HvRACB protein may thus support susceptibility. Two HvRACB-interacting barley proteins negatively regulate GTP-bound HvRACB. HvMAGAP1 is a microtubule associated ROP-GTPase activating protein (ROP-GAP) that likely stimulates GTP hydrolysis depending on the catalytic arginine-finger of its GAP domain (Hoefle et al., 2011). Barley ROP binding kinase1 (HvRBK1) is an active cytoplasmic receptor-like kinase, whose activity is stimulated by CA HvRACB *in vitro* and that directly binds to CA HvRACB *in planta* (Huesmann et al., 2012). HvRBK1 in turn interacts with components of an E3 ubiquitin ligase complex and controls protein abundance of activated HvRACB (Reiner et al., 2016).

Besides its role as an S factor, HvRACB appears to function in polar cell growth processes (Hoefle et al., 2011; Pathuri et al., 2008; Scheler et al., 2016). Other plant ROP GTPases act in plant immunity (Kawano et al., 2014). However, *HvRACB* does apparently not influence the ability of barley to express canonical PTI responses such as ROS generation and phosphorylation of mitogen-activated protein kinases (Scheler et al., 2016).

Here, we report on the HvRACB interacting *Bgh* <u>ROP</u>-<u>i</u>nteractive <u>p</u>eptide 1 (ROPIP1) that is encoded on the *Bgh* SINE-like retroposon Eg-R1. Our study suggests ROPIP1 to act as secreted intracellular virulence factor of *Bgh*.

## RESULTS

### ROPIP1 Is Encoded by the Retrotransposable Element Eg-R1 of *Bgh*

We performed yeast-two hybrid (Y2H) screens using the barley ROPs HvRACB (GenBank: AJ344223), CA HvRACB and CA HvRAC1 (GenBank: AJ518933) as baits against a cDNA library prepared from *Bgh*-infected barley leaves. Besides barley proteins (Hoefle et al., 2011; Huesmann et al., 2012), a *Bgh*-derived cDNA was repeatedly isolated. Sequencing of the respective plasmids isolated from yeast retrieved a poly-adenylated transcript and fragments of the same transcript that aligned to its 5’-region. Initial BLAST searches against the NCBI nucleotide database identified the transcript as the Non-LTR retroelement Eg-R1 (Wei et al., 1996) (GenBank: X86077.1) of *Bgh*. The Eg-R1 5’-sequence as obtained from the fragments in frame with the activation domain of the prey vector would give rise to a 74 amino acids peptide (see Fig. S1) which interacted with HvRACB, CA HvRACB and CA HvRAC1 in the bait vectors. Later on, we named this peptide <u>ROP</u>-<u>I</u>NTERACTIVE <u>P</u>EPTIDE 1 (ROPIP1) of *Bgh*.

A BLAST search of the Eg-R1 nucleotide sequence against the assembled *Bgh* reference genome (BGH DH14 Genome v3b (contigs); blugen.org) of race DH14 suggested more than 3000 genomic insertions of the Eg-R1 element and similar numbers in other fungal races (Hacquard et al., 2013). This number is likely underestimated as e.g. only half of the genome of *Bgh* race A6 could get assembled due to the high repeat content (Hacquard et al., 2013). We randomly selected 53 full-length Eg-R1 genomic insertions for inspection of the direct genomic environment. Interestingly, eight of the 53 insertions showed 5’-elongated ORFs (Table S1) including the 74 amino acids that had been isolated in the Y2H screening in frame with predicted signal peptides for secretion (SignalP 3.0 Server). This would extrapolate to hundreds of such genomic sequences given at least 3000 genomic insertions. 5’-RACE-PCR further confirmed (Table S2) the recently published Eg-R1 consensus sequence (Eg-R1_cons) (Amselem et al., 2015). BLAST searches of the ROPIP1 or the Eg-R1 nucleotide sequence against the NCBI nucleotide collection exclusively produced hits matching to the species *Blumeria graminis* possibly hinting to a specificity of the Eg-R1 element for powdery mildews of *Poaceae*. A highly similar retroelement, Bgt_RSX_Lie, was identified in the genome of the close *Bgh*-relative *Bgt* of wheat (Parlange et al., 2011).

Eg-R1 was originally described as repetitive element that shares some features with SINEs but which is also distinct from classical SINEs (Wei et al., 1996). SINEs typically share sequence similarities with tRNAs, 7SL RNA or 5S rRNA from which they may derive (Kramerov and Vassetzky, 2011). All these are transcribed by RNA pol III. As already reported by Wei and colleagues (1996) (Wei et al., 1996) Eg-R1 lacks A-box and B-box RNA pol III transcription initiation sites within its 5’-region. Furthermore, internal poly (T) stretches would act as RNA pol III termination signals such that a RNA pol III transcript would be truncated which renders transcription by RNA pol III very unlikely. Genomic insertions of Eg-R1 lacked genomic poly (A)-coding stretches at their 3’-ends but comprised a 5’-AAUAAA-3’ polyadenylation signal, which is obviously functional since Eg-R1 is expressed as polyadenylated RNA ((Wei et al., 1996) Fig. S1; see Fig. S2 for Eg-R1 architecture). This supports gene-like transcription of Eg-R1 by RNA pol II. The ROPIP1 nucleotide sequence was amplifiable from cDNA prepared from total RNA extracts as well as from poly (A) mRNA preparations of *Bgh*-inoculated barley leaves but not from the non-inoculated control (Fig. S3). Wei and colleagues (1996) detected Eg-R1 on a northern blot of poly (A) RNA (Wei et al., 1996). Expression of ROPIP1 and Eg-R1 was further supported by BLAST searches against expressed sequence tags of *Bgh* (BGH DH14 All ESTs database) of race DH14 (blugen.org) and RNAseq data of *Bgh* race A6 grown on the immunocompromised *Arabidopsis thaliana* (Hacquard et al., 2013). Genomic insertions of Eg-R1 were found located in close spatial vicinity to CSEPs, where Eg-R1 was suggested to mediate unequal crossing over events (Pedersen et al., 2012). This might be supported by our finding of truncated Eg-R1 genomic insertions not being reflected by preferential insertion of an Eg-R1 partial sequence, which could have arisen from e.g. incomplete insertion of the element (Fig. S4A, S4B). Eg-R1 is deposited at Repbase (Repbase Report 2011, Volume11, Issue 9, (Jurka et al., 2005)) as one member of an eight *Blumeria graminis* Non-LTR retrotransposons (BG_Non-LTRs)-comprising family, which were found to be conserved in their 5’-region (Fig. S4C, S4D). In summary, the ROPIP1 sequence was found encoded on Eg-R1, which likely is a member of a class of yet not well characterized, non-autonomous, RNA pol II-transcribed retroelements.

### ROPIP1 Interacts with Barley Susceptibility Factor HvRACB in Yeast

We next verified the ROPIP1-HvRACB protein interaction in yeast by independent targeted Y2H assays. Besides wild-type (WT) and CA HvRACB, the dominant negative mutant DN HvRACB and HvMAGAP1 (GenBank: AK371854) were additional included as bait proteins. Yeast colony growth of the prey-bait combinations ROPIP1-HvRACB and ROPIP1-CA HvRACB exceeded all other combinations on interaction-selective media (Fig. 1A). Weak background growth of the ROPIP1 prey was abolished when plating yeast on 2.5 mM 3-AT (Fig. 1B). No colony growth was observable when ROPIP1 was combined with either DN HvRACB or HvMAGAP1. There is no obvious ATG start at very 5’-end of the Eg-R1 nucleotide sequence we found in the Y2H screening. However, there is an ORF in the same reading frame of the ROPIP1 sequence, which translates into a peptide of 44 amino acids and which we refer to as ROPIP1-Cter (see Fig. S1). In order to delimit the HvRACB-interacting part, ROPIP1 was split into ROPIP1-Cter and the remaining N-terminus (ROPIP1-Nter). The fragments were tested against the same baits like ROPIP1 in targeted Y2H assays. ROPIP1-Cter in the prey vector did not show any background growth. ROPIP1-Cter interacted in yeast with CA HvRACB and HvRACB but not with DN HvRACB, which resulted in an identical pattern to ROPIP1 as prey. However, colonies grew less dense when compared to ROPIP1 (Fig. 1A). ROPIP1-Nter was not sufficient for interaction with any of the baits. Together this suggested that binding of ROPIP1 to HvRACB is largely mediated by ROPIP1-Cter. Secondary structure prediction for ROPIP1 proposed folding in α-helices and β-sheet structures (Fig. S4E, F).

**Fig. 1.**
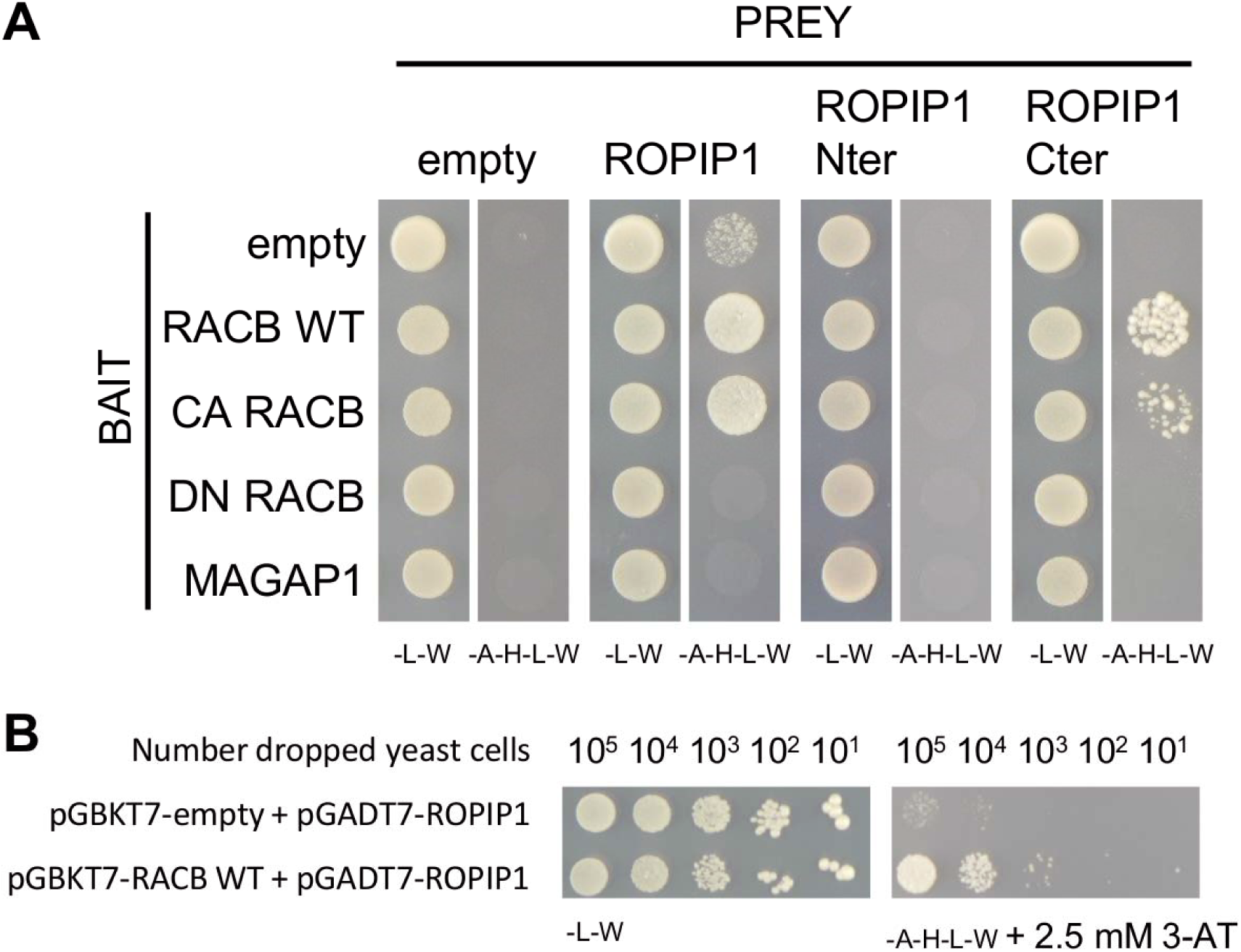
*Bgh* ROPIP1 and ROPIP1-Cter Interacted with Barley HvRACB and CA HvRACB in Yeast. (A) ROPIP1 of *Bgh* was tested as prey in targeted Y2H assays for interaction with the barley small GTPase HvRACB in three different variants: WT: wild-type protein, CA: constitutively activated mutant (HvRACB G15V), DN: dominant negative mutant (HvRACB T20N) and with the HvRACB-interacting protein HvMAGAP1. The ROPIP1 sequence was additionally split into its small inherent C-terminal ORF (ROPIP1-Cter) which was sufficient for protein interaction with HvRACB WT and CA HvRACB and the remaining N-terminal part (ROPIP1-Nter) which did not interact with the baits. 10^5^ cells of each combination were dropped in parallel on SD –Leu, -Trp (-L-W) as transformation control and on SD –Ade,-His, -Leu, -Trp (-A-H-L-W) selection medium. (B) Serial dilution of 10^5^ to 10 yeast cells transformed with pGADT7-ROPIP1 as prey vector and pGBKT7-HvRACB WT as bait vector or pGBKT7-empty as empty vector control. Left panel: transformation control medium (SD –L-W). Right panel: Selection medium (SD –A-H-L-W) supplemented with 2.5 mM 3-AT to increase selectivity.

### ROPIP1 Enhances Virulence of *Bgh*

As ROPIP1 interacted with the susceptibility factor HvRACB, we checked whether ROPIP1 can affect the susceptibility of barley against *Bgh*. Therefore, we transiently expressed ROPIP1 in barley epidermal cells by microprojectile bombardment prior to inoculation with *Bgh* conidial spores at 24 hours after transformation (hat) and microscopic analysis of fungal development at 48 hai. To express the full ROPIP1 sequence including the ROPIP1-Nter and ROPIP1-Cter *in planta*, we equipped the sequence with an additional ATG start codon at its very 5’-end (see Suplemental Table S1 and S2). Transformed cells were identified by co-bombarded GFP. Over-expression of ROPIP1 led to a significant increase (P ≤ 0.05, Student’s t-test) in susceptibility to fungal penetration of transformed barley leaf epidermal cells. This was evident from an enhanced frequency of attacked cells with fungal haustoria. Hence, ectopic expression of ROPIP1 promoted virulence of *Bgh* (Fig. 2A). The relative penetration rate increased thereby by around 40 percent. Ectopic over-expression of ROPIP1-Cter in barley epidermal cells had an effect comparable to albeit somewhat weaker than that of ROPIP1. This added to the view of ROPIP1-Cter being the part of ROPIP1 that promotes virulence of *Bgh*.

We went on with transient silencing of *ROPIP1*. No readily applicable transformation protocol is available yet for *Bgh*. However, ectopic expression of double-stranded RNAi constructs in barley epidermal cells proved to be a valuable tool for silencing *Bgh* transcripts in a process called host-induced gene silencing (HIGS) (Ahmed et al., 2015; Nowara et al., 2010; Pliego et al., 2013; Zhang et al., 2012). ROPIP1 was cloned as inverted repeat into the plant RNAi vector pIPKTA30N (Douchkov et al., 2005). Off targets prediction using the SI-FI software (Nowara et al., 2010) did neither reveal further targets in *Bgh* nor in barley. For the HIGS experiment, the transformed leaves were inoculated at 24 hat with *Bgh* conidia followed by microscopic analysis of fungal development at 48 hai. HIGS of ROPIP1 significantly (P ≤ 0.05, Student’s t-test) reduced the relative penetration rate of *Bgh* on transformed cells by 38% (Fig. 2B). We included a synthetic ROPIP1 RNAi-insensitive rescue construct in the experiment to assure that the observed drop in virulence of *Bgh* was due to posttranscriptional silencing of ROPIP1. The nucleotides at the wobble position in the codons of the ROPIP1 nucleotide sequence were exchanged to be most different to the codons used by *Bgh* in ROPIP1 but to be most similar to the codon usage of barley (Fig. S5C). The functionalities of the ROPIP1-RNAi and ROPIP1-RNAi rescue constructs were tested in advance by transient co-expression experiments and silencing GFP-ROPIP fusion constructs (Fig. S5A). Accordingly, ROPIP1-RNAi-rescue partially but significantly (P ≤ 0.05, Student’s t-test) rescued the ROPIP1-RNAi mediated decrease in fungal penetration success (Fig. 2B).

**Fig. 2.**
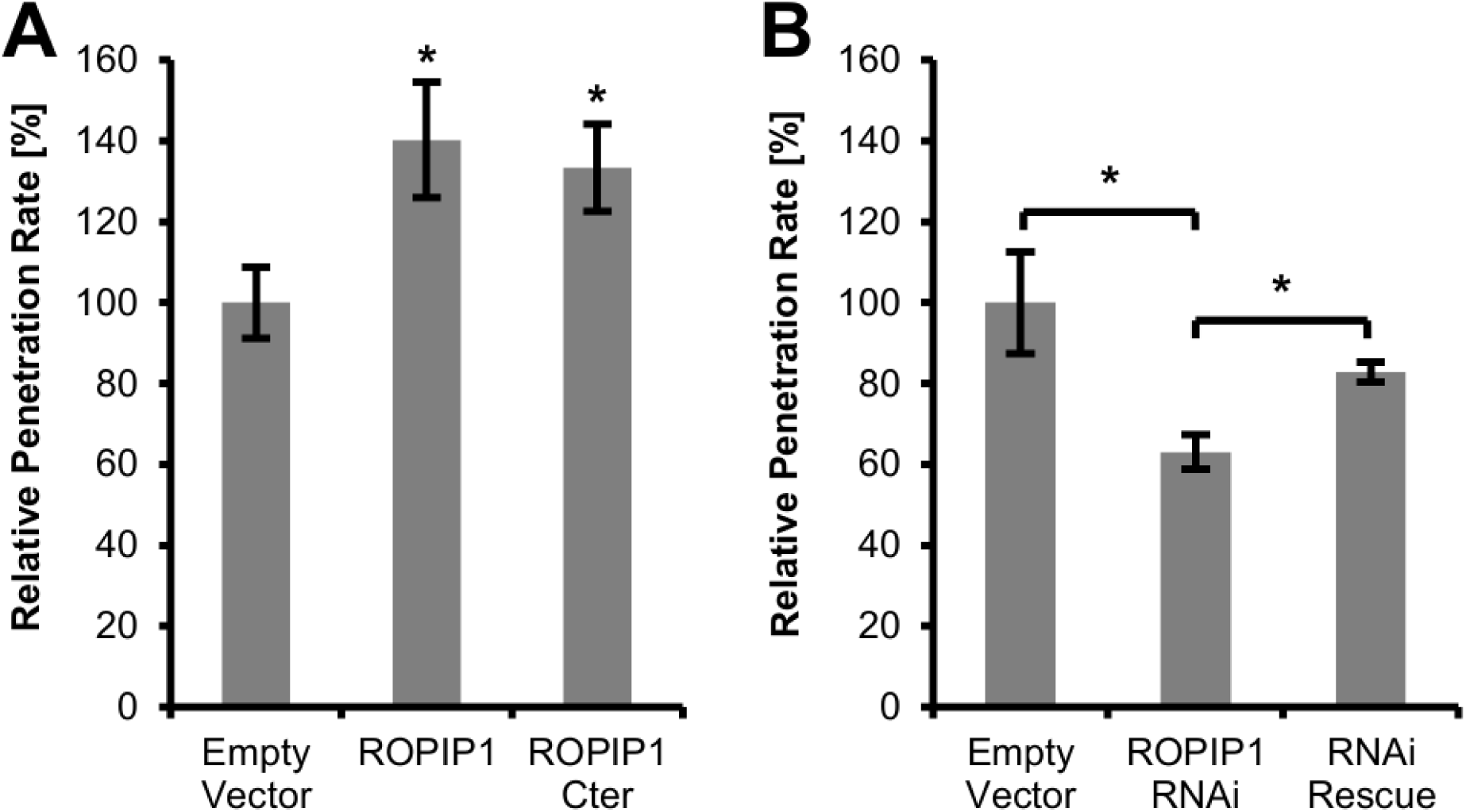
ROPIP1 Modulated Susceptibility of Barley Epidermal Cells towards *Bgh*. (A) Transient over-expression of ROPIP1 and ROPIP1-Cter in barley epidermal cells significantly increased the relative penetration rate of *Bgh* in comparison to the control. (B) Host-induced gene silencing (HIGS) of native *ROPIP1* by transient expression of ROPIP1 as dsRNA (ROPIP1-RNAi) in barley epidermal cells significantly decreased the relative penetration rate of *Bgh*. Co-expression of a ROPIP1 RNAi rescue construct (RNAi rescue) significantly complemented HIGS of the native *ROPIP1* transcript. Bars represent the mean values of six independent experiments in (A) and four independent experiments in (B). Error bars are ± SE. * P ≤ 0.05 (Student’s t-test).

### ROPIP1 Protein is Detecteble in *Bgh*-Infected Barley Leaf Protein Extracts

We next investigated whether a native ROPIP1 protein is detectable. A custom rabbit polyclonal antibody, α-ROPIP1, was raised against a synthesized epitope peptide derived from ROPIP1-Cter. The monospecific IgG fraction was purified to ≥ 95 % by affinity chromatography using the epitope peptide as antigen. Total protein extracts were prepared from heavily *Bgh* infected and non-inoculated barley primary leaves. A unique band in the protein extract of the *Bgh* inoculated sample was repeatedly observable in a series of western blots (Fig. 3A). The band was never seen in the protein extract prepared from non-inoculated samples. Recombinant, *E.coli*-expressed His-tagged ROPIP1 (recROPIP1) was run as positive control on the same gel and was detected by α-ROPIP1 (Fig. 3A). Further, α-ROPIP1 specifically detected recROPIP1 in crude cell lysates of *E. coli* cell cultures following induction of recombinant protein expression with isopropyl β-D-1-thiogalactopyranoside (IPTG). The identity of the signal was confirmed by, first, the absence of the band in the non-induced control; second, by probing aliquots of the same crude cell lysates with an independent α-His antibody, which resulted in an identical signal pattern (Fig. 3B).

**Fig. 3.**
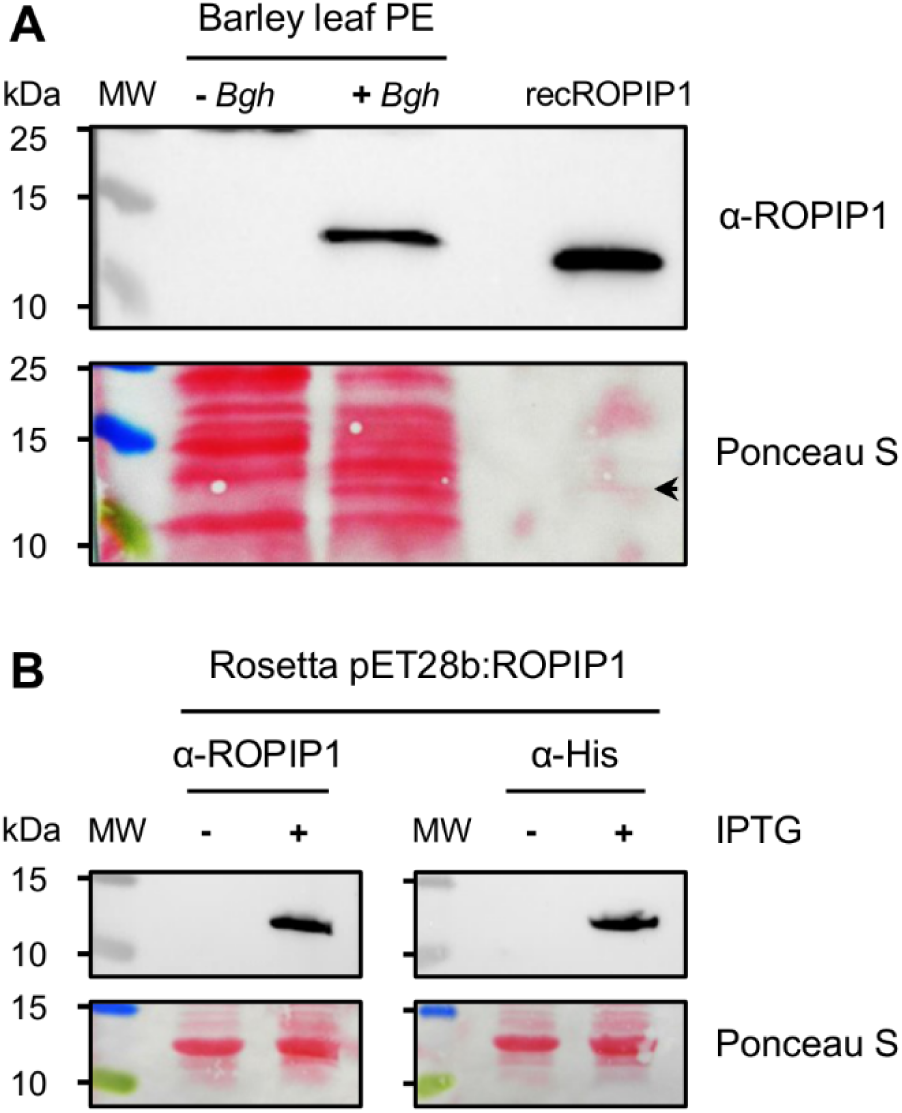
Western Blot of Barley Leaf Protein Extracts Using α-ROPIP1 Antibody. (A) Affinity purified anti-peptide antibody α-ROPIP1 was used as primary antibody in western blots of total protein extracts prepared from barley leaves inoculated (+*Bgh*) or non-inoculated (-*Bgh*) with *Bgh*. His-tag purified recombinant ROPIP1 (recROPIP1) was run as positive control on the same gel. RecROPIP1 and a protein exclusive to the +*Bgh* sample were labeled by α-ROPIP1. Several repetitions confirmed the signal in the +*Bgh* lane. (B) Controls for α-ROPIP1 specificity. *E.coli* Rosetta cells were transformed with the IPTG-inducible vector pET28b:ROPIP1. Crude cell lysates were prepared from small scale cell cultures with (+) or without (-) IPTG-induction. Recombinant His-tagged ROPIP1 was detected by α-ROPIP1 in the IPTG-induced sample (+) but not in the non-induced control (-). The usage of α-His antibody in aliquots of the same samples validated the identity of the signal. The experiment was repeated twice with identical results. Ponceau S: loading and protein transfer control. The arrowhead points to a faint band in the recROPIP1 lane in (A). MW: Molecular weight protein ladder. PE: Protein extract.

### TEM Localizes ROPIP1 in *Bgh* Structures and in the Host Cell Cytoplasm

Next, we analyzed the localization of the protein labeled by α-ROPIP1 *in situ*. We used immunogold labeling and transmission electron microscopy (TEM). Ultrathin cuts of heavily *Bgh*-infected (3 days after inoculation, dai) barley primary leaves were incubated with α-ROPIP1 or an unspecific antibody as primary antibodies. Primary antibodies were detected by anti-rabbit secondary antibodies conjugated to 10 nm gold particles.

Fungal intra- and extracellular structures, the extracellular space, the cell wall and the barley epidermal cell interior were almost free from gold particles in the unspecific antibody control (Fig. 4A and detail in 4B). By contrast, gold particles labeled fungal and host cell structures upon usage of α-ROPIP1 as primary antibody. In a barley epidermal cell, showing a host cell wall apposition (CWA, also called papilla), gold particles were found in the epicuticular fungal appressorium, inside the host cell wall and the host CWA (Fig. 4C and detail in 4D). Gold particles appeared to spread from the tip of the appressorium but were almost absent from the extracellular space and the host cell vacuole. Hence, α-ROPIP1 obviously targeted a secreted fungal protein. In a penetrated barley epidermal cell, where *Bgh* established a fungal haustorium, gold particles were located in the fungal haustorium as well as in the host cytoplasm (Fig. 4E and detail in 4F) but not in the host vacuole showing that epitopes were not displaced during sample preparation. Therefore, the α-ROPIP1 labelled protein apparently was able to translocate from the fungus into the cytoplasm of barley epidermal host cells. Almost no gold particles were detectable in mesophyll cells of *Bgh*-infected barley leaves. Very few gold particles were occasionally observed in plastids (Fig. S6).

In sum, immunogold labelling with α-ROPIP1 detected a secreted *Bgh* protein that translocated from the fungus into barley epidermal cells, where it could interact with HvRACB.

**Fig. 4.**
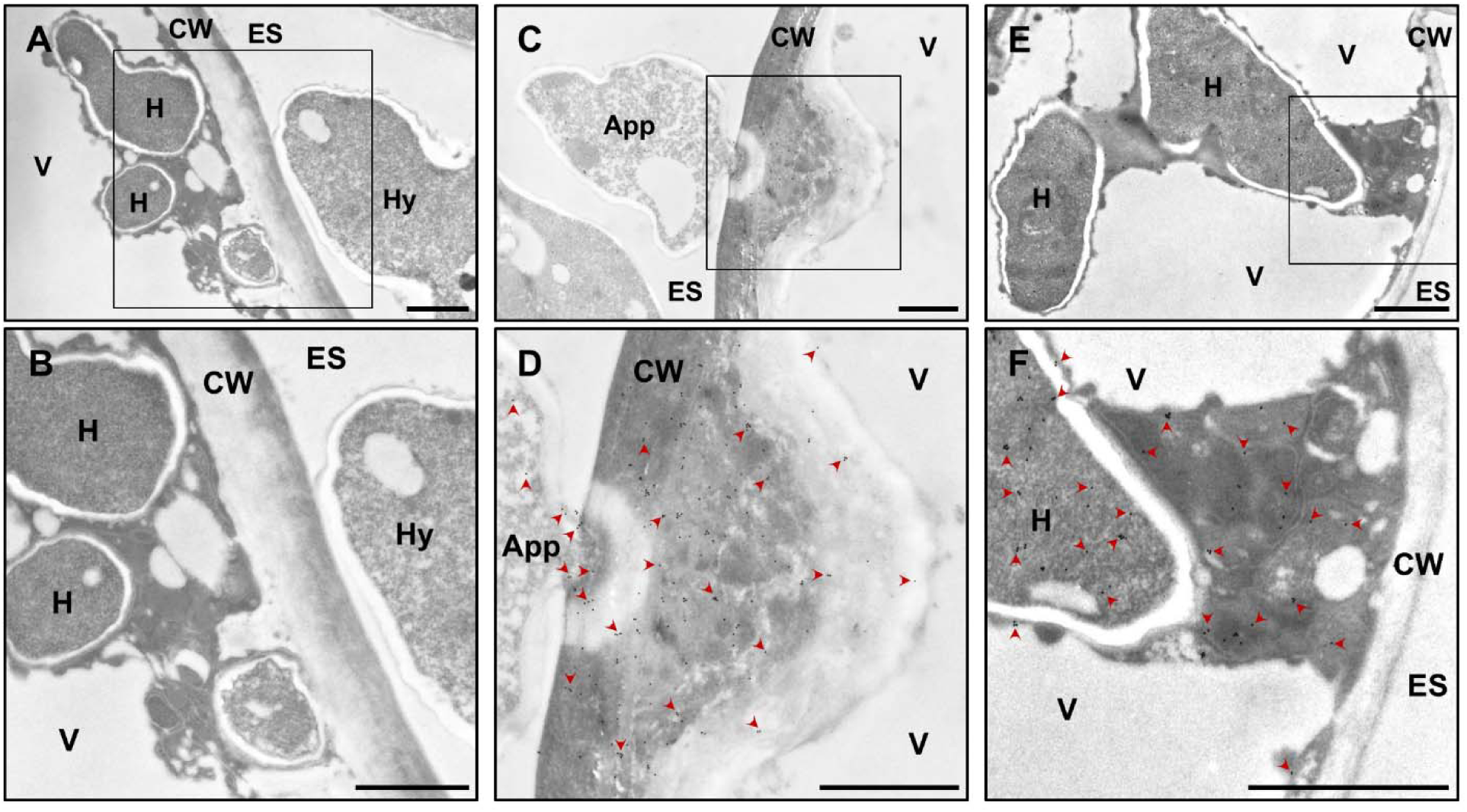
Immunogold labeling of α-ROPIP1 in *Bgh*-Challenged Barley Leaves. Transmission electron micrographs of ultrathin sections of *Bgh*-infected barley epidermal cells 3 dai showing gold particles bound to α-ROPIP1. (A, B) Negative control of infected cells treated with a non-specific antibody. Gold particles were absent in the susceptible barley epidermal cell containing intracellular fungal haustorial protrusions (H) and the extracellular *Bgh* hypha (Hy). (C, D) Gold particles bound to α-ROPIP1 were observed inside a *Bgh* appressorium (App), the barley epidermal cell wall (CW) and papilla but were absent from the extracellular space (ES) and the host cell vacuole (V). (E, F) Gold particles were found in the lumen of finger-like *Bgh* haustorial protrusions inside barley epidermal cells as well as the host cell cytoplasm but were almost absent from the host cell vacuole (V), the CW and the ES. Arrowheads in (D) and (F) point to selected gold particles. Scale bars are 1µm.

### HvRACB Binding HvMAGAP1 Recruits ROPIP1 to Microtubules

With ROPIP1 being a potential intracellular effector of *Bgh* we went on with live cell imaging of GFP-tagged ROPIP1 by confocal laser scanning microscopy. Transient expression of GFP-ROPIP1 in barley epidermal cells did not show a distinct subcellular localization of ROPIP1. GFP-ROPIP1 labeled the cytoplasm and the nucleus (Fig. 5A). This was in line with the ROPIP1 sequence not showing any predictable cellular localization signatures or protein domains. As HvRACB-interacting proteins associate with microtubules or function in regulation of MT network stability, we expressed GFP-ROPIP1 together with the putative HvRACB-regulator HvMAGAP1 that has a unique localization at MTs (Hoefle et al., 2011). Although ROPIP1 did not interact with HvMAGAP1 in yeast (Fig. 1), GFP-ROPIP1 was recruited to MTs under co-expression of red fluorescing RFP-HvMAGAP1 (Fig. 5B). The C-terminus of HvMAGAP1 (HvMAGAP1-Cter) mediates MT association of HvMAGAP1 but does not interact with HvRACB because it lacks the ROP-interacting CRIB motif and the GAP domains (Hoefle et al., 2011). Quantification subcellular fluorescence of GFP-ROPIP1 at 12 – 24 hat revealed that full length RFP-HvMAGAP1 recruited GFP-ROPIP1 to MTs whereas RFP-HvMAGAP1-Cter hardly co-localized with GFP-ROPIP1 at MTs (P ≤ 0.001, Χ^2^, Fig. 5C, 5D). Instead, GFP-ROPIP1 labeled the cytoplasm as did soluble GFP upon co-expression of RFP-HvMAGAP1 or RFP-HvMAGAP1-Cter (Fig. 5C, 5D). Hence, GFP-ROPIP1 localization at cortical MTs depended on RFP-HvMAGAP1 with its corresponding HvRACB-binding domains.

**Fig. 5.**
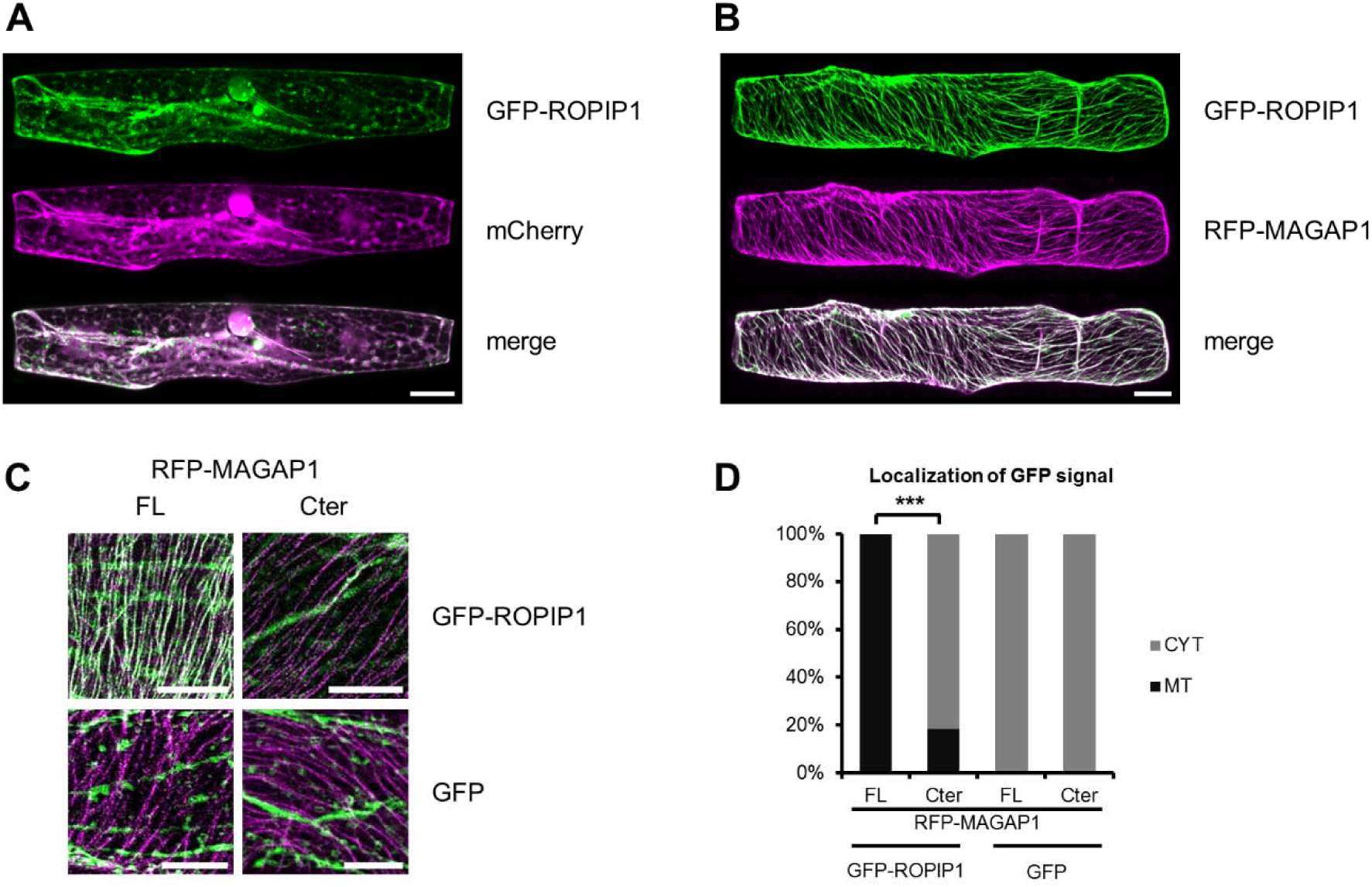
Recruitment of GFP-ROPIP1 to Cortical Microtubules by RFP-HvMAGAP1. Barley leaf epidermal cells were transiently transformed by particle bombardment and imaged with confocal laser scanning microscopy as sequential whole cell scans in 2 µm increments at 12 – 24 hat. (A) Whole cell projection showing cytoplasmic and unspecific subcellular localization of GFP-ROPIP1. Co-localization with cytoplasmic and nucleoplasmic mCherry fluorescence is indicated by white pixels in the merge picture. The observation was consistently repeatable in more than three experiments. (B) Recruitment of GFP-ROPIP1 to cortical microtubules (MT) upon co-expression of MT-associated RFP-HvMAGAP1. White pixels in the merge picture indicate co-localization. A maximum projection of 20 optical sections in 2 µm increments is shown. The observation was consistently repeatable in more than three experiments. (C) Visualization of co-expressed fusion protein combinations used for quantitative analysis. C-ter: Truncation of HvMAGAP1 to the MT-associated C-terminus (HvMAGAP1-Cter). FL: full-length HvMAGAP1. 10 optical sections of the upper cell cortex were merged for the pictures. (D) Quantification of the combinations shown in C. Bars are frequencies of cells with GFP fluorescence being located at MTs or in the cytoplasm only (CYT) derived from three independent experiments. The respective absolute numbers of the categories were compared in a Χ²-test. RFP-HvMAGAP1-Cter highly significantly reduced MT-association of GFP-ROPIP1 (*** p ≤ 0.001, n = 61, 60, 53, 57 cells from left to right). Scale bars in A, B, C are 20 µm.

### ROPIP1 and CA HvRACB Interact *in planta* and Can Co-Localize With HvMAGAP1

To support that ROPIP1 can interact with activated HvRACB in *planta*, we performed ratiometric bimolecular fluorescence complementation (BiFC, Figs. 6A-C). ROPIP1-YFP^N^ was transiently co-expressed with either YFP^C^-CA HvRACB or YFP^C^-DN HvRACB, RFP-HvMAGAP1-R185G, a mutant lacking the catalytic arginine finger of GAP domains (Hoefle et al., 2011) and CFP. The RFP-HvMAGAP1 R185G mutant was chosen as its co-expression with ROPIP1 was seen to influence the organization of the cortical MT network less than co-expression of RFP-HvMAGAP1, which destabilized MTs in presence of ROPIP1. However, RFP-HvMAGAP1 R185G interacts with CA HvRACB in *planta* (Hoefle et al., 2011), and GFP-ROPIP1 was recruited to MTs by RFP-HvMAGAP1-R185G (suppl. Fig. S7). Ratiometric measurement of YFP versus CFP signals showed fluorescence complementation of ROPIP1-YFP^N^ with YFP^C^-CA HvRACB but only weakly with YFP^C^-DN HvRACB or YFP^C^-HvMAGAP1 (Fig. 6A, suppl. Fig. S8). The mean YFP/CFP ratio of YFP^C^-CA HvRACB co-expressing cells was significantly different from that in YFP^C^-DN HvRACB-co-expressing or YFP^C^-HvMAGAP1 cells (P ≤ 0.01 or 0.001, repsctively Student’s t-test; Fig. 6C, suppl. Fig. S8). The BiFC signal of ROPIP1-YFP^N^ and YFP^C^-CA HvRACB was predominantly observed at the cell periphery and as filamentous strings at the cell cortex, likely representing cortical MTs (Fig. 6B, 6D). Localization at the cell periphery is indicative for the plasma membrane, as activated HvRACB is partially plasma membrane-associated (Schultheiss et al., 2003). This supported a direct protein-protein interaction of ROPIP1-YFP^N^ and YFP^C^-CA HvRACB but not with YFP^C^-DN HvRACB or YFP^C^-HvMAGAP1 *in planta*. Localization of the BiFC signal at filamentous structures suggested that ROPIP1, activated HvRACB and HvMAGAP1 are simultaneously present at MTs, when co-expressed. This was supported by co-localization of GFP-ROPIP1, CFP-CA HvRACB and RFP-HvMAGAP1 at both MTs and the cell periphery (Fig. 6D).

**Fig. 6.**
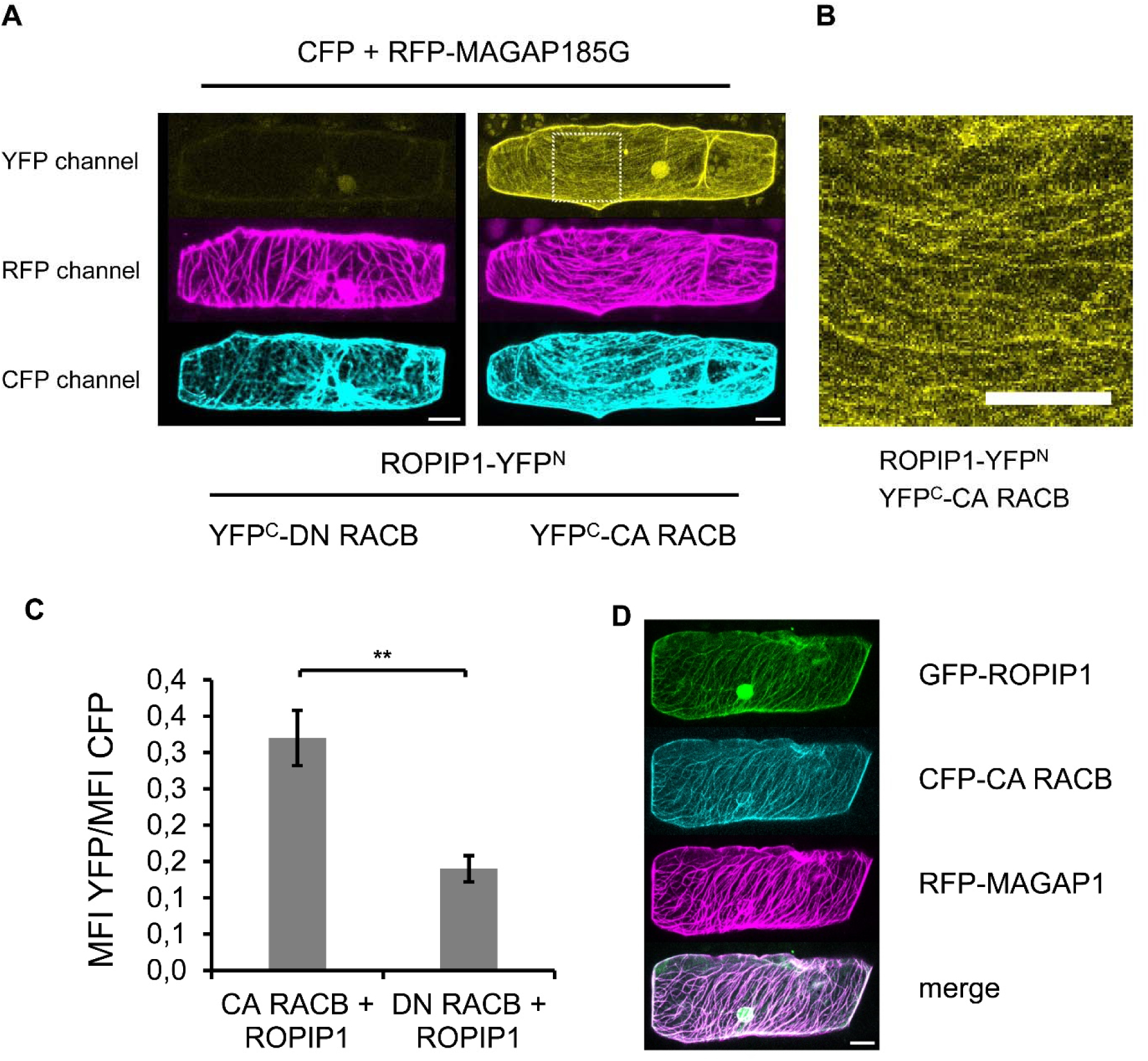
Split YFP Complementation of ROPIP1-YFP^N^ and YFP^C^-CA HvRACB *in planta*. (A) ROPIP1-YFP^N^ was transiently co-expressed with DN (Adamakis et al.) or CA (right) YFP^C^-HvRACB, the inactive RFP-HvMAGAP1-R185G mutant and CFP as transformation marker in barley leaf epidermal cells. Confocal laser scanning microscopy whole cell maximum projections are shown. (B) Detail picture of the ROPIP1-YFP^N^ and YFP^C^-CA HvRACB co-expressing cell from A (dashed square). A maximum projection of 10 optical sections à 2 µm from the upper cell cortex is shown. Scale bars in A and B are 20 µm. (C) Ratiometric measurement of YFP fluorescence complementation. ROPIP1-YFP^N^ was transiently co-expressed with YFP^C^-CA HvRACB or YFP^C^- DN HvRACB and YFP signals were normalized to signals from co-expressed CFP. Error bars are ± S.E. Two-sided Student’s t-test (**; P ≤ 0.01). (G) Co-Expression of GFP-ROPIP1, CFP-CA HvRACB and RFP-HvMAGAP1. Transformed cells were imaged as whole cell scans by confocal laser scanning microscopy at 48 hat. GFP-ROPIP1, CFP-CA HvRACB and RFP-HvMAGAP1 showed similar localization at the cell periphery and at microtubules. Scale bar is 20 µm.

### ROPIP1 Causes Microtubule Network Destabilization

HvMAGAP1 is located on MTs and supports focusing of MTs towards the site of attempted pathogen entry (Hoefle et al., 2011). We hence asked whether the recruitment of ROPIPI1 to MTs by HvMAGAP1 could influence MT organization. RFP-HvMAGAP1 was co-bombarded into barley epidermal cells with either GFP*ROPIP1 or GFP as control. We assessed MT organization on the basis of the three categories: intact MT network, disordered MT network, or fragmented MT network (Fig. 7A). Co-expression of GFP-ROPIP1 together with RFP-HvMAGAP1 led to a highly significant change (P ≤ 0.001, Χ^2^ test) in the distribution of the three categories when compared to control cells (Fig. 7B). The relative amount of category 3 cells exhibiting a fragmented MT network tripled from 15% in control cells to around 45% in cells co-expressing GFP-ROPIP1 and RFP-HvMAGAP1.

**Fig. 7.**
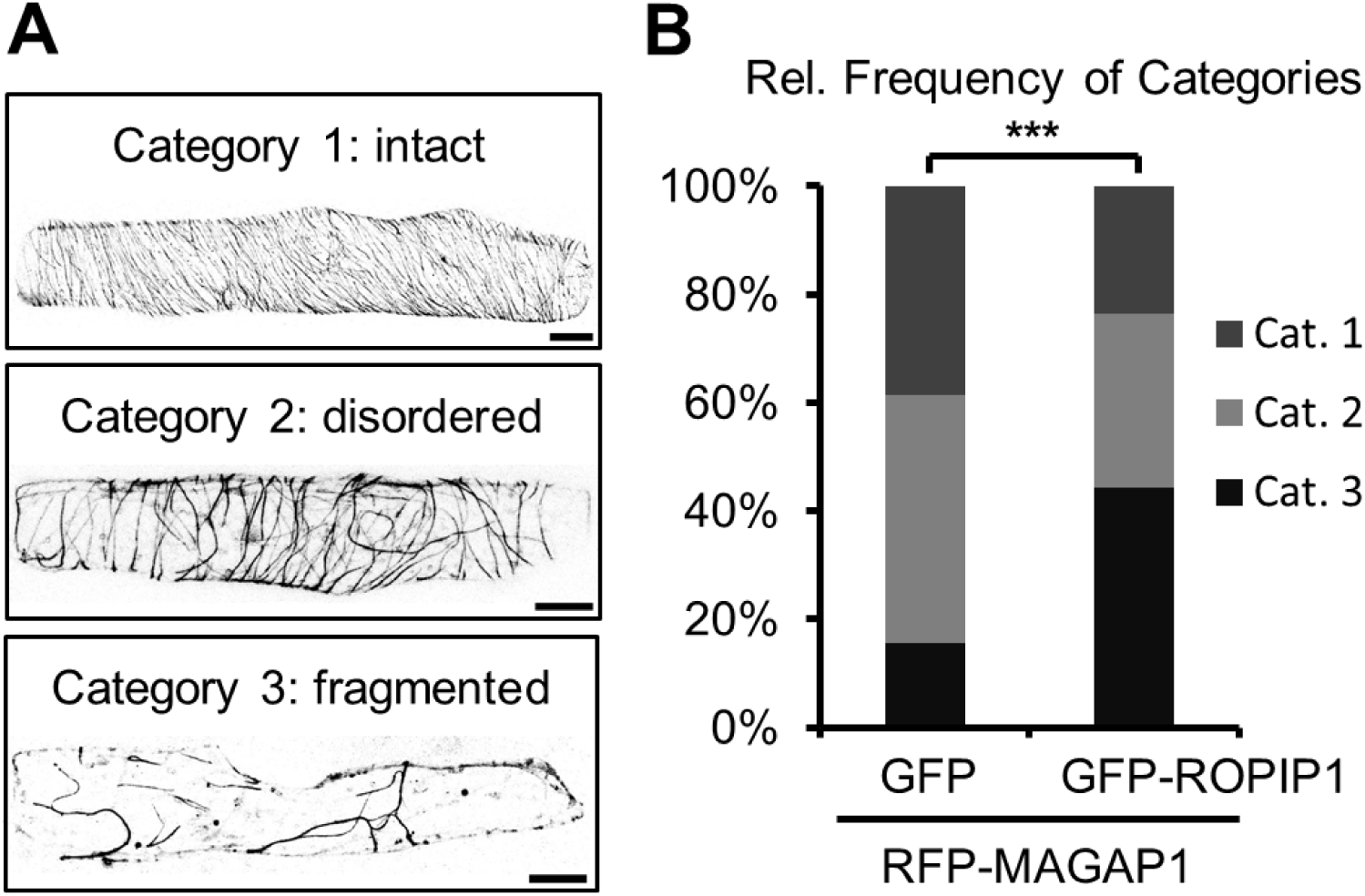
Co-Expression of GFP-ROPIP1 and RFP-HvMAGAP1 Enhanced Microtubule Network Disorganization. (A) Example micrographs illustrating three distinguished categories of MT network organization in barley epidermal cells. Confocal laser scanning microscopy whole cell projections of barley epidermal cells transiently co-expressing GFP-ROPIP1 and RFP-HvMAGAP1 are shown in grey scale. Scale bars are 20 µm. (B) Mean relative frequencies of the categories at 12 – 24 hat. The absolute numbers of cells per category of n = 145 GFP and n = 132 GFP-ROPIP1 cells each co-transformed with RFP-HvMAGAP1 obtained from four independent repetitions were compared in a Χ^2^-test (*** p ≤ 0.001; Χ^2^ = 27.92; df = 2). Cells of category 3 exhibiting a heavily disorganized microtubule network tripled from 15.5 % in the GFP control to 44.3 % in cells expressing GFP-ROPIP1.

## DISCUSSION

We identified the retroelement-encoded peptide ROPIP1 of *Bgh* that shows the potential to interact with the barley susceptibility factor HvRACB and to promote fungal penetration success on barley. Some *B. graminis* effectors have recently been characterized. Direct interaction with potential host target proteins has been reported for CSEP0055 that interacts with the barley pathogenesis-related protein PR17c (Zhang et al., 2012) and for CSEP0105 and CSEP0162 that interact the small heat shock proteins 16.9 and 17.5 (Ahmed et al., 2015). Additionally, a combination of protein pull down and Y2H experiments, CSEP0064 interacted with a glutathione-S-transferase, a malate dehydrogenase and a pathogenesis-related-5 protein isoform (Pennington et al., 2016). Some *B. graminis* effector candidates do not possess N-terminal signal peptides for secretion though they are thought to act intracellularly. This is the case for the class of EKA effectors (Ridout et al., 2006) and Candidate Effector Proteins (CEPs) of the wheat powdery mildew *Bgt* (Wicker et al., 2013). EKA effector genes are evolutionary and transcriptionally linked with autonomous Non-LTR retroelements (Ridout et al., 2006; Sacristan et al., 2009), whereas CSEPs and flanked by non-autonomous non-LTRs like *Eg-R1* and *Egh24* (Pedersen et al., 2012). Recent findings suggest that Avr_a10_ and Avr_k1_ evolved from insertions of premature stop codons in LINE ORF1 protein (ORF1p), which subsequently underwent positive selection (Amselem et al., 2015). This further supports potential neo-functionalization of *Bgh* retroelements as a genetic resource for the evolution of novel effector proteins.

The ROPIP1 sequence is distributed in the genome of *Bgh* by Eg-R1 but does not encode an N-terminal signal-peptide. The N-terminal ROPIP1 sequence part is not equipped with a canonical start codon on Eg-R1 whereas ROPIP1-Cter could be translated from an internal ATG. This raises the future question whether there might be a gain of function through formation of chimeric ORFs or whether the C-terminal peptide ORF ROPIP1-Cter represents the actual effector. Inspection of the barley genome readily revealed the presence of several chimeric ORFs which encoded extended stretches of amino acids and partially N-terminal signal-peptides that are in frame with ROPIP1. Due to the repetitive nature of ROPIP1 and the consequent presence of thousands of copies, the genomic origin of the detected polyadenylated RNAs and the corresponding protein remain unresolved and need further investigations. Our western blot experiment suggested that a ROPIP1-related sequence indeed gets translated into protein, because the used antibody against a ROPIP1-Cter peptide detected both recombinant ROPIP1 and a protein which was only present in *Bgh*-infected leaves. Immunogold-labeling and TEM further supported that this protein is secreted by the fungus and translocated into the host cell. Protein signal appeared in the infecting fungus and infected cells but did neither appear in uninfected barley nor mesophyll of infected barley. Hence, a host-translocated and intracellularly acting protein of *Bgh* was detected by the α-ROPIP1 antibody. Since α-ROPIP1 also detected recombinant ROPIP1 expressed from *E.coli*, we suggest that a ROPIP1 or a ROPIP1-related protein was detected in the fungus and the host cell cytoplasm. Further, ROPIP1 interacted with the barley susceptibility factor HvRACB in yeast and *in planta*. Hence, the barley small GTPase HvRACB likely is the host target of an ROPIP1 effector. Some first insights into a possible mode of action of ROPIP1 were gained. GFP-ROPIP1 co-located with HvRACB-interacting RFP-HvMAGAP1 at cortical microtubules in barley epidermal cells. Transient over-expression of GFP-ROPIP1 together with RFP-HvMAGAP1 promoted the breakdown of the cortical MT array. A possible protein complex containing ROPIP1, activated HvRACB and its regulator HvMAGAP1 was indicated by localization of a BiFC complex of the ROPIP1-CA HvRACB at filamentous strings at the cell periphery, which appear to represented cortical MTs, due to recruitment of ROPIP1 by HvMAGAP1 to MTs. MTs are involved in penetration resistance to powdery mildew fungi and RACB and RACB-like ROP GTPases are key regulators of MT (Hoefle et al., 2011; Huesmann et al., 2012; Kobayashi et al., 1997). Hence, the potential of ROPIP1 to destabilize cortical MTs suggests the possibility that ROPIP1 acts by manipulation of host MT arrays. This could either inhibit polarized cell wall-associated defense mechanisms or more directly facilitate fungal invasion and membrane delivery for formation of the extrahaustorial membrane and matrix (Dörmann et al., 2014).

### ROPIP1 – a *Bgh* Effector of Retroelement Origin

ROPIP1 does not fit predefined categories or definitions of secreted effector proteins of filamentous plant pathogens or prokaryotic or eukaryotic pathogens in general. However, that does not exclude ROPIP1 from being an effector. There are recent published examples that expand the current model of plant pathogen effectors beyond strict definitions. The effectors PsIsc1 and VdIsc1 of the oomycete *Phytophthora sojae* (*P. sojae*) and the phylogenetically distinct true fungus *Verticillium dahliae*, respectively attenuate the PTI response by misdirecting the synthesis of the plant defense hormone salicylic acid. Both proteins do not encode N-terminal signal peptides for secretion but PsIsc1 can functionally replace the N-terminal signal peptide and the RXLR-dEER host translocation motif of the effector Avr1b of *P. sojae* (Liu et al., 2014). This adds to the assumption that there should be an additional secretion pathway besides the conventional co-translational loading into the endomembrane route or a process of cytoplasma exchange with host cells in filamentous plant pathogens possibly involving exosome release from multivesicular bodies (Micali et al., 2011).

ROPIP1 constitutes an unconventional effector candidate whose evolution was possibly supported by the high repeat content of the *Bgh* genome. It would be of great interest to learn whether there are comparable proteins getting expressed in other species. The finding of long intergenic non-coding (linc) RNAs being translated in the human proteome provoked the view that presumably non-coding RNAs constitute an evolutionary playground (Wilhelm et al., 2014). Similarly, ribosome profiling identified 5’-regions of about 10–100 codons of yeast long non-coding RNAs to be bound by ribosomes, which suggests their translation (Smith et al., 2014). By looking at ROPIP1 we are possibly observing the neo-functionalization of a non-coding retroelements into a new effector gene. The nature of the Eg-R1 element has to be characterized further as it shares some SINE properties but different from SINE elements, it is obviously transcribed by RNA pol II ((Wei et al., 1996) and this study). A recent report proposed a similar class of SINE-like non-autonomous but RNA pol II transcribed SINEU retroelements from crocodilians to represent a new class of retroelements or at least a new class of SINEs (Kojima, 2015). Apart from that, there is a great lack in knowledge about this type of non-autonomous retroelements. The 5’-region of SINEU-element family members is derived from small nuclear U1 and U2 genes that obviously were recurrently added to the 5’-end of SINEU ancestor elements (Kojima, 2015). Interestingly, the Eg-R1 sequence part following the 3’-end of the ROPIP1 part showed some relatedness to classical SINE element architecture (Fig. S2).

TE-rich genomic islands are suggested as origin regions of effector diversification in the genomes of *P. infestans* and *Leptosphaeria maculans brassicae* (Grandaubert et al., 2014; Raffaele et al., 2010). These plasticity islands are typically gene-poor but enriched in effector genes and are thought to speed up evolution by providing a highly dynamic source of genetic variation including asexual sequence variation (Raffaele et al., 2010). The whole *Bgh* genome is primarily composed of TEs with genes being interspersed in small clusters. It is one of the biggest ascomycete genomes possibly due to the absence of a TE spread controlling repeat-induced point mutation (RIP) mechanism (Spanu et al., 2010). The high repeat content may give myriads of options for non-allelic recombination making the genome very dynamic. The SINE-like Eg-R1 retroelement, the SINE Egh24 and the high copy repeat element AJ002007.1 were found to closely (~0.5 to ~1kb) surround CSEPs and might provide sites for unequal crossing-over events. CSEP diversification is thought to be driven by frequent gene-duplications (Pedersen et al., 2012). Eg-R1 can be found located in close spatial vicinity to likely secreted classical effector proteins. The current knowledge is too sparse to clearly draw a conclusion on the evolution of a possibly virulence promoting sequence being dispersed throughout the genome by an SINE-like retroelement. Anyhow, the experimental data suggest an effector function of a ROPIP1 sequence containing protein. It further appears possible that ROPIP1 gained an N-terminal signal peptide by insertional formation of chimeric ORFs like those exemplary identified in this study (Table S1). Even if this should not be the case, ROPIP1 or ROPIP1-Cter yielded scores for predicted non-classical protein secretion comparable to those of PsIsc1 and VdIsc1 using the SecretomeP 2.0 server (Bendtsen et al., 2004) in analogy to (Liu et al., 2014). Predicted protein folding (Fig. S4) but absence of predictable functional domains in ROPIP1 is typical as many effector proteins represent novel folds which implies the possibility that they are not derived from sequence variation of preexisting genes. Further, gene losses of the primary and secondary metabolism of *B. graminis*, likely due to high retrotransposon activity, reflect its extreme obligate biotrophic lifestyle (Spanu et al., 2010; Wicker et al., 2013) which greatly enhances selective pressure. In a genome with a gene set being reduced to the necessary minimum, non-gene transcripts may gain novel functionalities in virulence and in general.

## METHODS

### Plant Growth and Pathogen Infection

Barley (*Hordeum vulgare* L.) cultivar ‘Golden Promise’ was grown at 18 °C, 60 % relative humidity in a photo period of 16 h and a photon flux of 150 µmol s^-1^m^-2^. *Blumeria graminis* (DC) Speer f.sp. *hordei* Em. Marchal, race A6 (Wiberg, 1974) was propagated on barley cultivar ‘Golden Promise’ under the same conditions. For protein extraction, 7 d old barley plants were inoculated with >150 conidia/mm² and let grown until 10 dai. First leaves were inoculated with ~ 150 conidia/mm² for RT-PCR and harvested at the indicated time points or were inoculated with ~300 conidia/mm² and let grown until 3 dai for immunogoldlabeling and TEM. Transiently transformed detached 7 d old primary leaves kept on 0.5 % water-agar were inoculated with ~ 150 conidia/mm² at 24 hat.

### Targeted Y2H

ROPIP1 was identified by DNA sequencing of positive prey clones from a yeast two-hybrid screen using HvRACB, CA HvRACB and CA HvRAC1 as baits against a cDNA library prepared from *Bgh*-infected barley leaves, published in (Hoefle et al., 2011). For targeted Y2H assays, yeast strain AH109 MATa was co-transformed with pGBKT7 bait plasmids and pGADT7 prey plasmids following the small-scale LiAc yeast transformation procedure (Clontech, Heidelberg, Germany).

ROPIP1-Nter was PCR-amplified from pGADT7-ROPIP1 using primer V42A_SmaI_F and R_V42A_Nter_BamHI (Supplemental Table S3) and SmaI/BamHI cloned into pGADT7. ROPIP1-Cter was PCR-amplified from C-ROPIP1 using primer F_V42ACter_Sma and R_V42ACter_Bam and SmaI/BamHI cloned into pGADT7. Cloning of barley ROP proteins into pGBKT7 vector is described in (Schultheiss et al., 2008). Transformed cells were selected on SD medium lacking Leu and Trp (-L-W), resuspended in ultrapure water and spotted on SD –L-W and on interaction selective SD medium lacking Ade, His, Leu and Trp (-A-H-L-W). 3-amino-1,2,4-triazole (3-AT) was optionally added in concentrations from 0.5–2.5 mM to the SD –A-H-L-W medium to increase selectivity.

### Transient Transformation of Barley Leaf Epidermal Cells

Primary leaves of 7 d old barley plants were cut and placed on solid 0.5 % water-agar. Plasmids were coated to 1.0 µm gold-particles (BioRad) and bombarded into barley epidermal cells using the PDS-1000/He (Bio-Rad) system as described earlier (Douchkov et al., 2005; Eichmann et al., 2010).

### Transient Over-Expression and HIGS

For transient over-expression ROPIP1, respectively ROPIP1-Cter were PCR-amplified from cDNA using 5’-oligos V20A,V42ABamH1fwd, respectively V42A,V20BBamH1kurz and 3’-oligo V42A,V20Brev, A/T-cloned into pGEM-T (Promega) and BamHI/SalI subcloned into the pUC18-based pGY1-plant expression vector (Trujillo et al., 2006). An 5’-ATG start codon for ROPIP1 *in planta* expression was introduced into the ROPIP1 sequence by the 5’-oligo V20A,V42ABamH1fwd. Detached barley primary leaves were co-bombarded with 0.5 µg/shot pGY1-GFP for transformation control and 1.0 µg/shot of pGY1-ROPIP1 or pGY1-ROPIP1-Cter or pGY1 empty vector. Microscopic evaluation of haustoria formation in GFP-fluorescing cells was at 48 hai. The relative penetration efficiencies were calculated by dividing the number of transformed cells with haustoria by the sum of susceptible plus resistant (attacked by *Bgh* but stopped) transformed cells of each combination. In each combination and repetition at least 50 cell autonomous interactions scored. The relative penetration rate was calculated by forming the quotient of the penetration efficiency of each sample divided by the penetration efficiency of the control. The variation of the control samples was calculated by dividing the penetration efficiency from each repetition by the arithmetic mean of all penetration efficiencies of the control samples. The arithmetic means calculated from the relative penetration efficiencies of the test samples were pairwise compared to the arithmetic means of the relative penetrations efficiencies of the control in a two-sided Student’s t-test. For transient HIGS, ROPIP1 was PCR-amplified from cDNA using the 5’-oligo V20A,V42ABamH1fwd and the 3’-oligo V42A,V20Brev and blunt-ligated into the Gateway entry vector pIPKTA38. ROPIP1 was then recombined as inverted repeat into the Gateway destination vector pIPKTA30N by standard Gateway LR reaction (Douchkov et al., 2005). The synthetic ROPIP1-RNAi rescue (Eurofins MWG Operon) was designed by replacing the original codons by the most different but not rare barley codons (see Supplementary Fig. 3C) as described by (Nowara et al., 2010). The codon usage frequencies were obtained from the Codon Usage Database (http://www.kazusa.or.jp/codon/). ROPIP1-RNAi-rescue was BamHI/SalI subcloned from the delivered pEX-A2 plasmid into the pGY1 plant expression vector. GFP was cloned in frame with ROPIP1-RNAi-rescue into the BamHI cleavage site resulting in pGY1-GFP-ROPIP1-RNAi-rescue. For the HIGS experiment 1.0 µg/shot of pIPKTA30N-ROPIP1, or empty pIPKTA30N (control) plus either 1.0 µg/shot pGY1-ROPIP1-RNAi-rescue or empty pGY1 and 0.5 µg /shot pGY1-GFP each were bombarded into barley epidermal leaf cells. Assessment of fungal development on GFP-expressing cells took place at 48 hat as described above for the over-expression experiment.

### Western Blot

Total protein extracts from heavily *Bgh*-infected barley primary leaves or mock-treated control leaves were prepared using the Plant Total Protein Extraction kit (Sigma-Aldrich) following the manufacturer’s instructions. Around 200 mg of liquid N 2-ground barley leaf powder was used for 250 µl of Protein Extraction Reagent Type 4. The protein concentration was determined by a Bradford assay. 50–100 µg of total protein per lane were separated by SDS-PAGE on hand-cast mini-gels (15 % resolving gel, 4 % stacking gel) using the Mini-PROTEAN Tetra Cell (Bio-Rad) in the Laemmli (Laemmli, 1970) buffer system. 200 V were applied for up to 45 minutes. Separated proteins were blotted onto 0.2 µm nitrocellulose membranes using a Fastblot B43 (Biometra) semi-dry blot system. A current of 5 mA/cm² was applied for 25 minutes. Successful protein transfer was checked by Ponceau S staining. Nitrocellulose membranes were destained by 2 rounds of washing in 1x PBS for 10 min, before blocking in 5.0 % non-fat dry milk in PBS for 1 h at RT. The blot was incubated with diluted primary antibodies (total barley protein extracts: 1:100, recombinant *E. coli* crude lysates: 1:10.000) in blocking buffer over-night at 4 °C. After three rounds of washing in PBS-T for each 15 min, blots were incubated with anti-rabbit-Hrp (Sigma-Aldrich) secondary antibodies diluted 1:80.000 in blocking buffer for 2 h at RT and washed again for three rounds. The SuperSignal West Femto Maximum Sensitivity Substrate (Thermo Scientific) was used as ECL substrate. Chemiluminescence was documented with a Fusion-SL4 system operated with FusionCapt Advance Solo 4 (version 16.06) software. The custom antipeptide antibody α-ROPIP1 (Pineda Antibody Service; Berlin, Germany) was raised against the synthesized peptide NH2-IPSRLRDLYRLHF-COOH in rabbits in a 145 d custom-controlled immunization protocol and purified to ≥ 95 % by affinity chromatography.

### Heterologous Expression of Recombinant ROPIP1

ROPIP1 was PCR amplified from plasmid using primer B8B,V21B_BamH1fwd and V42A,V20Bsalrev (Table S3) and BamHI/SalI cloned into the pET28b(+) vector. The pET28b-ROPIP1-6His plasmid was further digested with NdeI/BamHI to excise additional ATG start codons in the MCS. Sticky ends were blunted and the plasmid religated. The resulting pET28b-6His-ROPIP1-6His plasmid was transformed into chemically competent Rosetta (DE3) *E.coli* cells.

For crude-cell lysate preparation 50 ml of LB Kan (50 µg/ml kanamycine) were inoculated with a 1:100-diluted over-night culture. Small-scale cultures were grown until they reached an OD_600_ of 0.8 to 1.0. Non-induced aliquots were taken. Recombinant protein expression was induced by addition of IPTG to a final concentration of 1 mM. Induced and parallel non-induced cultures were grown at 37 °C for an additional 1–3 hours. Crude cell lysates were prepared by resuspending bacterial pellets in 100 µl Lysis Buffer (50 mM NaH2PO4-H2O, 300 mM NaCl, 10% glycerol, 1% Triton X-100, 1mg/ml Lysozyme, pH 8.0) per 1 ml of culture volume and incubation on ice for 30 min. Three rounds of ultrasonic bath incubation for 10 sec with placing on ice in-between followed. Viscosity of lysates was reduced by addition of 50 u Benzonase (Merck Millipore) per 1 ml culture volume and a further incubation on ice for 15 min. Up to 10 µl of heat-denatured crude lysate were loaded per lane onto SDS-PAGE gels. Non-induced control samples and IPTG-induced samples were run as duplicates on the same gels followed by western blotting. Afterwards, one half of the nitrocellulose membrane was incubated with α-ROPIP1 as primary antibody and the duplicate halve was incubated with anti-His-Hrp (Carl Roth). RecROPIP1 was purified with the Protino Ni-TED 2000 packed columns kit (Macherey Nagel) following the batch gravity-flow purification protocol under native conditions (User Manual, version Rev.04, protocol 5.5).

### Immunocytohistochemical Detection of α-ROPIP1

Sample preparation for TEM and immunogold labeling were performed according to a modified version described previously (Redkar et al., 2015). Briefly, samples were fixed with 2.5% (w/v) paraformaldehyde and 0.5% (v/v) glutaraldehyde in 0.06 M Sørensen phosphate buffer, then rinsed in buffer, dehydrated in acetone and embedded in LR-White resin (London Resin). Immunogold labeling of α-ROPIP1 was performed on ultrathin sections with an automated immunogold labeling system (Leica EM IGL, Leica Microsystems). The sections were blocked for 20 min with 2% (w/v) BSA (Sigma-Aldrich) in PBS, pH 7.2, and then treated with the primary antibody α-ROPIP1 against ROPIP1 for 90 min diluted 1:100 in PBS containing 1%(w/v) BSA. After sections were washed twice for 5 min with PBS containing 1% (w/v) BSA they were treated with a 10-nm gold-conjugated secondary antibody (goat anti-rabbit IgG, British BioCell International) diluted 1:100 in PBS containing 1% (w/v) BSA for 90 min. After a short wash in PBS (3 × 5 min) labeled grids were post-stained with 2% uranyl-acetate aqueous solution for 15 s and then investigated with a Philips CM10 TEM. The ideal dilutions and incubation times of the primary and secondary antibody were determined in preliminary studies by evaluating the labeling density after a series of labeling experiments. The final dilutions used in this study showed a minimum background labeling outside the sample with a maximum specific labeling in the sample. Various negative controls were performed to confirm the specificity of the immunocytohistochemical approach. Gold particles were absent on sections when a) no primary antibody, b) a nonspecific secondary antibody (goat anti-mouse IgG), and c) preimmune serum instead of the primary antibody was used.

### Live Cell Imaging

Transiently transformed barley epidermal leaf cells expressing fluorophore fusion proteins were imaged with a Leica TCS SP5 confocal laser scanning microscope using standard wavelengths for excitation and emission. Barley epidermal cells were scanned as z-stacks in 2 µm increments in sequential scan mode. Maximum projections were exported from the Leica LAS AF software (version 2.5.1) in jpeg or tiff format.

### Quantification of GFP-ROPIP1 Microtubule Localization and Destruction

GFP was cloned in frame with ROPIP1 into the 5’-BamHI restriction site of pGY1-ROPIP1 to gain pGY1-GFP-ROPIP1. The cloning of pGY1-RFP-HvMAGAP1 and variants is described in (Hoefle et al., 2011). Barley epidermal cells were transiently transformed with 0.5 µg/shot pGY1-GFP or 0.75 µg/shot pGY1-GFP-ROPIP1 plus 1.0 µg/shot pGY1-RFP-HvMAGAP1 or 1.0 µg/shot pGY1-RFP-HvMAGAP1-Cter and imaged as whole cell scans with 2 µm increments at 12–24 hat. For quantification of MT localization of GFP-ROPIP1 cells were categorized into GFP-signal present at MTs or absent from MTs. The numbers of categorized cells were compared between cells co-expressing RFP-HvMAGAP1 or RFP-HvMAGAP1-Cter together with GFP-ROPIP1 in a Χ² test with df = 1. For quantification of microtubule network organization maximum projections were categorized into intact, disordered or fragmented MTs. The distribution of the absolute cell numbers per category was compared between cells co-expressing GFP or GFP-ROPIP1 along with RFP-HvMAGAP1 in a Χ^2^ test with df = 2.

### Bimolecular Fluorescence Complementation

ROPIP1 was PCR-amplified from plasmid using 5’-oligo V20A,V42ABamH1fwd and 3’-oligo V42A,V20Bsalrev and BamHI/SalI cloned into the MCS of pUC-SPYNE (Walter et al., 2004) which translated into ROPIP-YFP^N^. The cloning of pUC-SPYCE-CA HvRACB, respectively pUC-SPYCE-DN HvRACB both translating into an N-terminal fusion of YFP^C^ to CA/DN HvRACB is described in (Schultheiss et al., 2008). Barley leaf epidermal cells (7d old) were transiently co-transformed with 0.75 µg/shot pUC-SPYNE-ROPIP1 plus 0.75 µg/shot pUC-SPYCE-CA HvRACB, respectively pUC-SPYCE-DN HvRACB, 0.5 µg/shot pGY1-CFP and 1.0 µg/shot pGY1-RFP-HvMAGAP1-R185G. Transformed cells were identified by CFP-fluorescence and imaged by confocal laser scanning microscopy at 36 hat. Each fluorophore was excited and detected in an individual scan by sequentially scanning between frames. All hardware and software settings were kept identical for all cells and repetitions. The BiFC signal was analyzed in a quantitative manner using maximum projections of transformed cells and the Leica LAS AF (version 2.5.1.6757) ‘Quantify’ tool. The first region of interest (ROI 1) was put at the cell periphery of the transformed cell. The second, copy-pasted, ROI 2 was placed into the surrounding background close to the cell. The mean values of fluorescence intensity of the ROIs (mean fluorescence intensity, MFI) of the YFP and the CFP detector were read out from the quantification reports. The background fluorescence MFI (ROI 2) was subtracted from ROI1. The corrected MFI of the YFP detector was divided by the corrected MFI of the CFP detector. The obtained YFP/CFP MFI ratios of YFP^C^-CA HvRACB and YFP^C^-DN HvRACB co-expressing cells were compared in a two-sided Student’s t-test. The corrected CFP MFIs were also compared in a two-sided Student’s t-test and did not differ.

### 5’-RACE-PCR

The Dynabeads mRNA Direct Kit (Thermo Scientific) was used according to the manufacturer’s instruction for isolation of poly (A) RNA from *Bgh*-infected barley primary leaves. After DNAseI digestion, the isolation process was repeated. 0.5–1.0 µg of poly (A) RNA were reverse-transcribed into first-strand cDNA following the instructions of the 5’/3’ RACE kit, 2^nd^ Generation, version 12 (Roche) and using the oligo TW42A_R as cDNA synthesis primer (Table S3). The resulting dA-tailed cDNA was used as template for PCR-amplification using Phusion High-Fidelity DNA Polymerase (Thermo Scientific) and V42A-SP2 as gene-specific primer. V42A-SP3 was used as nested gene specific primer in a second PCR run. PCR products were gel purified, A-tailed, cloned into pGEM-T (Promega) and sequenced.

### Semi-Quantitative RT-PCR

*Bgh*-inoculated and mock-treated barley primary leaves (7 d old) were cut and immediately frozen in liquid N2. Total RNA was prepared (Chomczynski and Sacchi, 1987). Total RNA was NaAc/ethanol precipitated to achieve greater purity and digested with DNAseI (Thermo Scientific). First strand cDNA was synthesized with RevertAid Reverse Transcriptase (Thermo Scientific) using Oligo (dT)15 primer (Promega). Barley *Ubiquitin Conjugating Enzyme 2* (*HvUBC2*, AY220735.1) gene was amplified using the oligo pair HvUBC2_fwd and HvUBC2_rev. Barley *Basic PR-1-Type Pathogenesis Related Protein* (*HvPR1b*, X74940.1) gene was amplified using the oligo pair T-PR1b/5´-2 and T-PR1b/3´-2. *Bgh Tub2 Gene For Beta Tubulin* (*Bgh tub2*, AJ313149) gene was amplified using the oligo pair *Bgh*_beta-tub_F and *Bgh*_beta-tub_R. *Bgh ROPIP1* transcript was amplified using the oligo pair V42fwd and V42rev.

## AUTHOR CONTRIBUTIONS

M.N. designed and carried out experiments, evaluated data and wrote the manuscript. C.M. and B.Z. performed experiments and evaluated data. R.H. developed the research questions, designed the study, supervised experimental work and wrote the manuscript.

## ACKNOWLEDGMENTS

The authors appreciate initial lab work on ROPIP1 by Jutta Preuss (Chair of Phytopathology, Technical University of Munich, Germany). We are grateful to Ruth Eichmann (University of Warwick, United Kingdom) for technical advice.

---

**Fig. S1.**
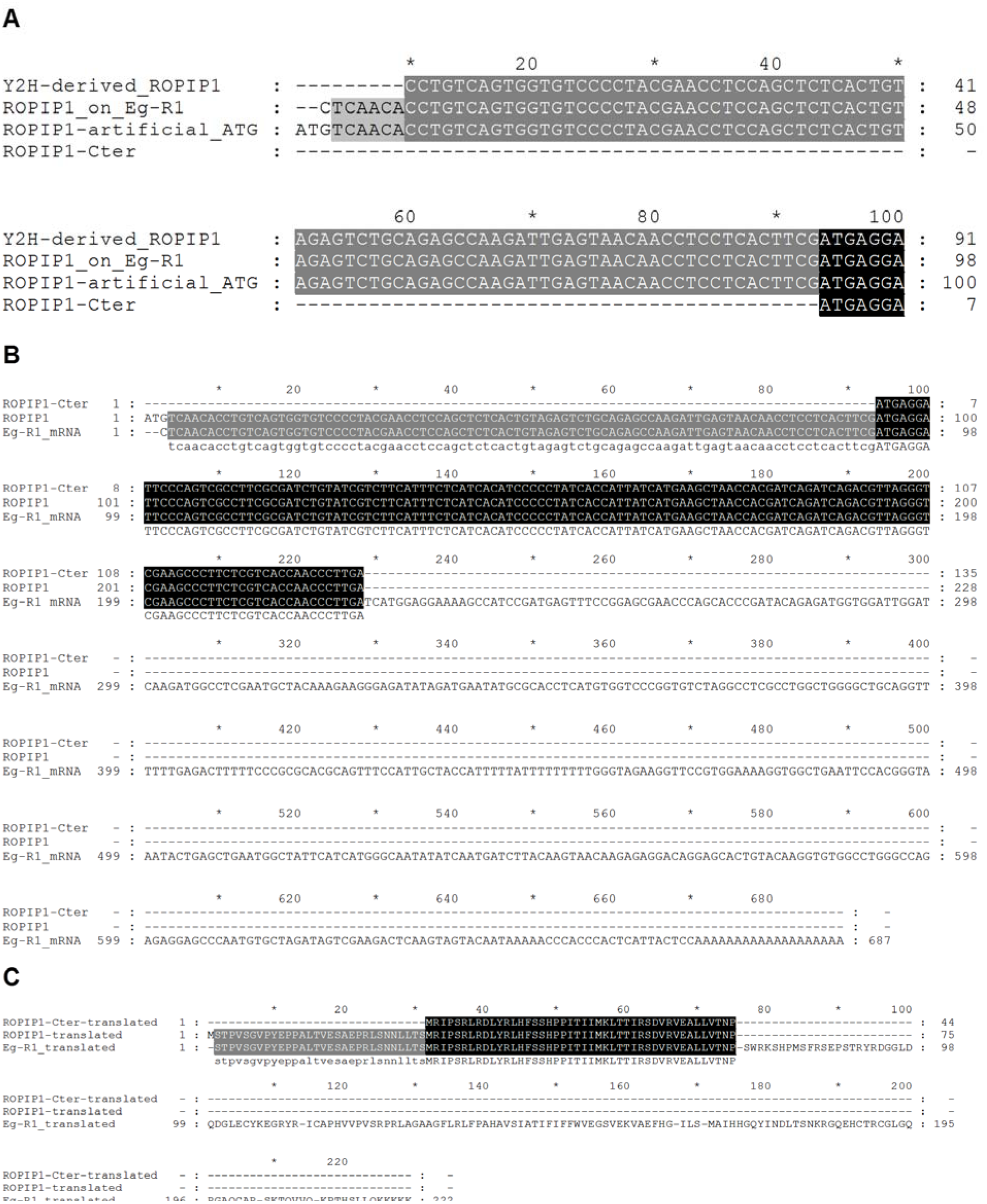
Sequence Alignments of Eg-R1, ROPIP1 and ROPIP1-Cter. (A) Nucleotide sequence alignment of the 5’-ends of the Y2H-derived ROPIP1 sequence, the annotated EgR1 (X86077.1) sequence, the ROPIP1 sequence equipped with an artificial 5’-ATG and the start of the intrinsic ORF represented by ROPIP1-Cter. (B) Nucleotide sequence alignment of Eg-R1, ROPIP1 and ROPIP1-Cter. (C) Amino acid sequence alignment of Eg-R1, ROPIP1 and ROPIP1-Cter.

**Fig. S2.**
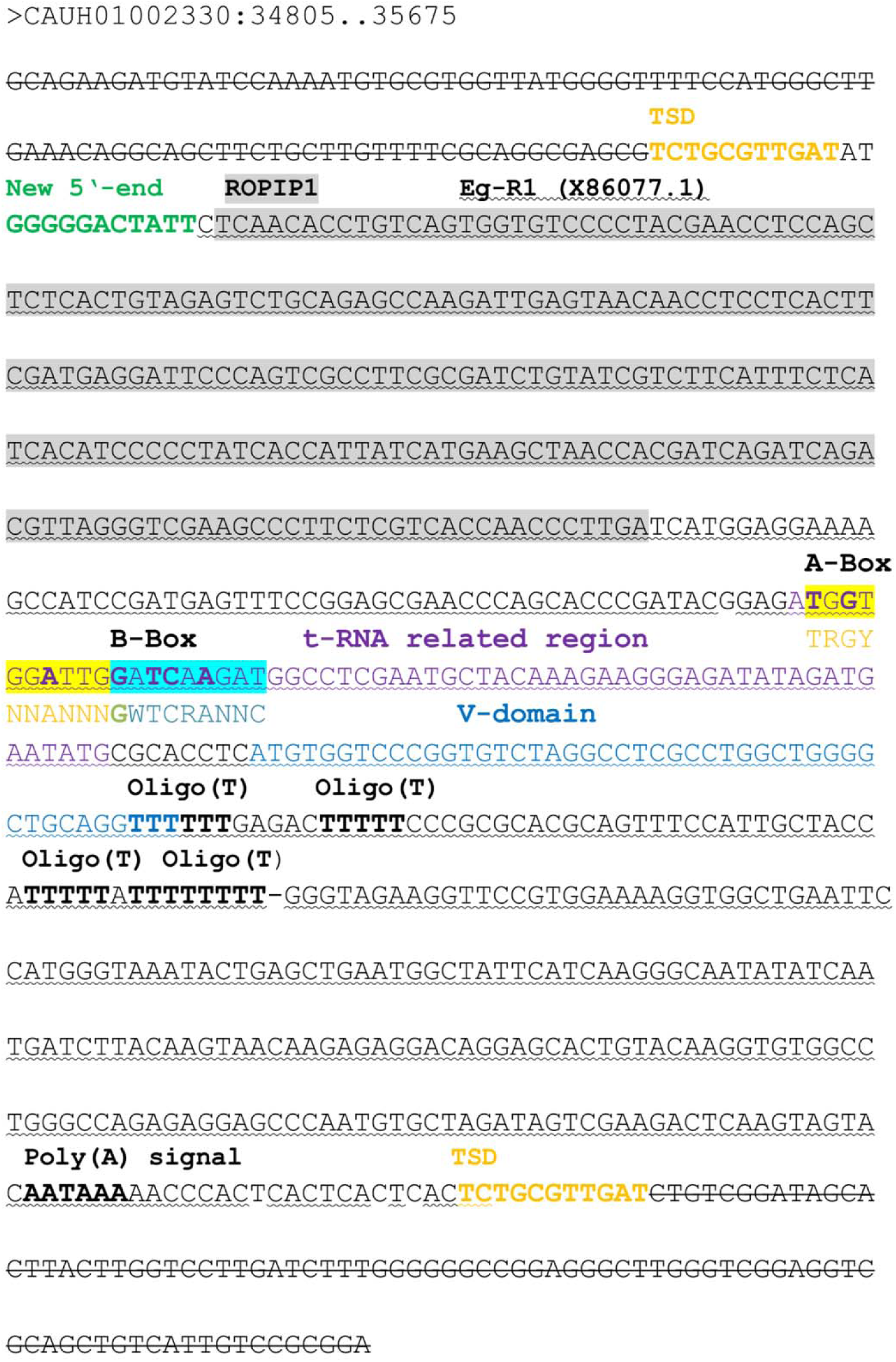
Exemplary Genomic Insertion and Hypothetical Architecture of the Eg-R1 Retroelement. This exemplary genomic insertion of a full-length Eg-R1 retroposon was extracted from contig CAUH010023330 at nucleotide position 34805–35675 of the *Bgh* genome (blugen.org). The surrounding genomic sequence is stroked-through. Likely retrotransposition-derived target site duplications (TSDs) are highlighted in orange. The 5’-end extension of 11 bp missing in the Eg-R1 annotation (X86077.1) is highlighted in green. Nucleotides identical to Eg-R1 are underlined. Nucleotides identical to ROPIP1 are shaded in grey. A putative SINE-like region following the ROPIP1 sequence part was identified by following the protocol for SINE analysis of SINEBase (Vassetzky and Kramerov, 2012). A putative tRNA-Gln (tdbD00008587|Homo_sapiens|9606|Gln|CTG; sines.eimb.ru) - related sequence region is highlighted in violet. Putative A- and B-Box-like sequences are shaded in yellow and turquoise. A-Box and B-Box pol III promoter consensus sequences determined for *Saccharomycetes* of the *Ascomycota* phylum (Marck et al., 2006) are given below the respective nucleotides. Matching nucleotides are in bold-type. Note that the putative A- and B-Box-like sequences are overlapping at one nucleotide highlighted in green. A region similar to the vertebrate SINE body part V-domain is highlighted in blue. Oligo(T)-stretches that would terminate pol III transcription are in bold-type. A likely functional polyadenylation (Poly(A)) signal is indicated in bold-type near the 3’-end of Eg-R1.

**Fig. S3.**
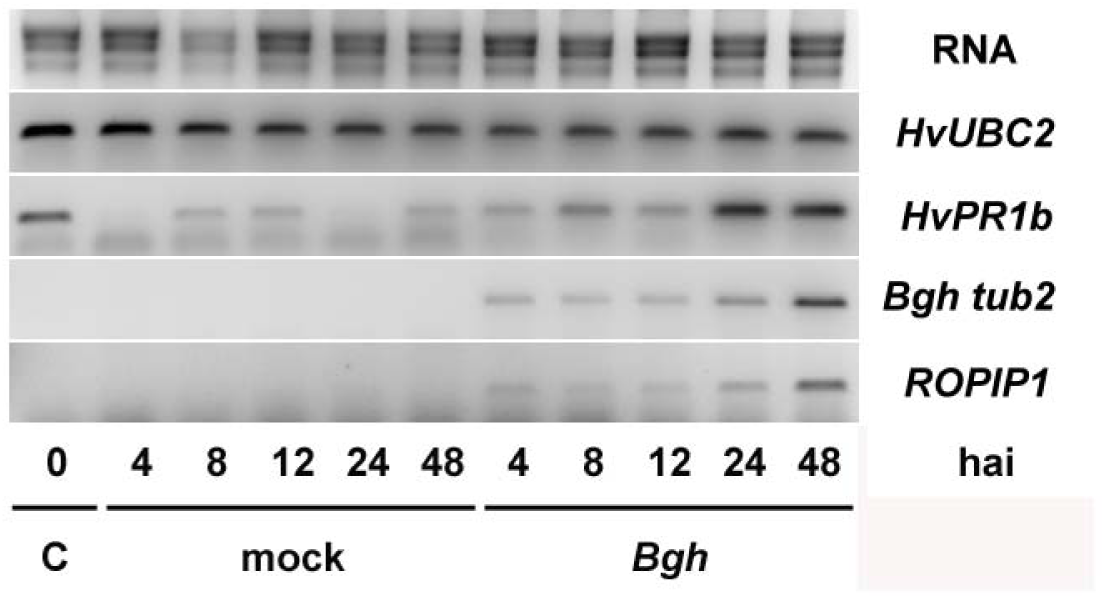
Semi-Quantitative Reverse Transcription PCR. 7d old barley primary leaves were inoculated with *Bgh* or mock treated. Leaves were cut and frozen in liquid N2 at the indicated time points. Total RNA was extracted, DNAse I-digested and reverse transcribed into cDNA. Barley *Ubiquitin Conjugating Enzyme 2 (HvUBC2*, AY220735.1) was amplified as control for cDNA quantity. Successful inoculation was checked by induction of the barley *Basic PR-1-Type Pathogenesis Related Protein (HvPR1b*, X74940.1) gene. *Blumeria graminis* f.sp. *hordei Tub2 Gene For Beta Tubulin (Bgh tub2*, AJ313149) was amplified to monitor the development of fungal biomass. The *ROPIP1* transcript appeared not to be induced after pathogen challenge. Being part of the SINE-like Eg-R1 retrotransposon the amplified ROPIP1 sequence is indistinguishable from Eg-R1. The experiment was repeated with similar results.

**Fig. S4.**
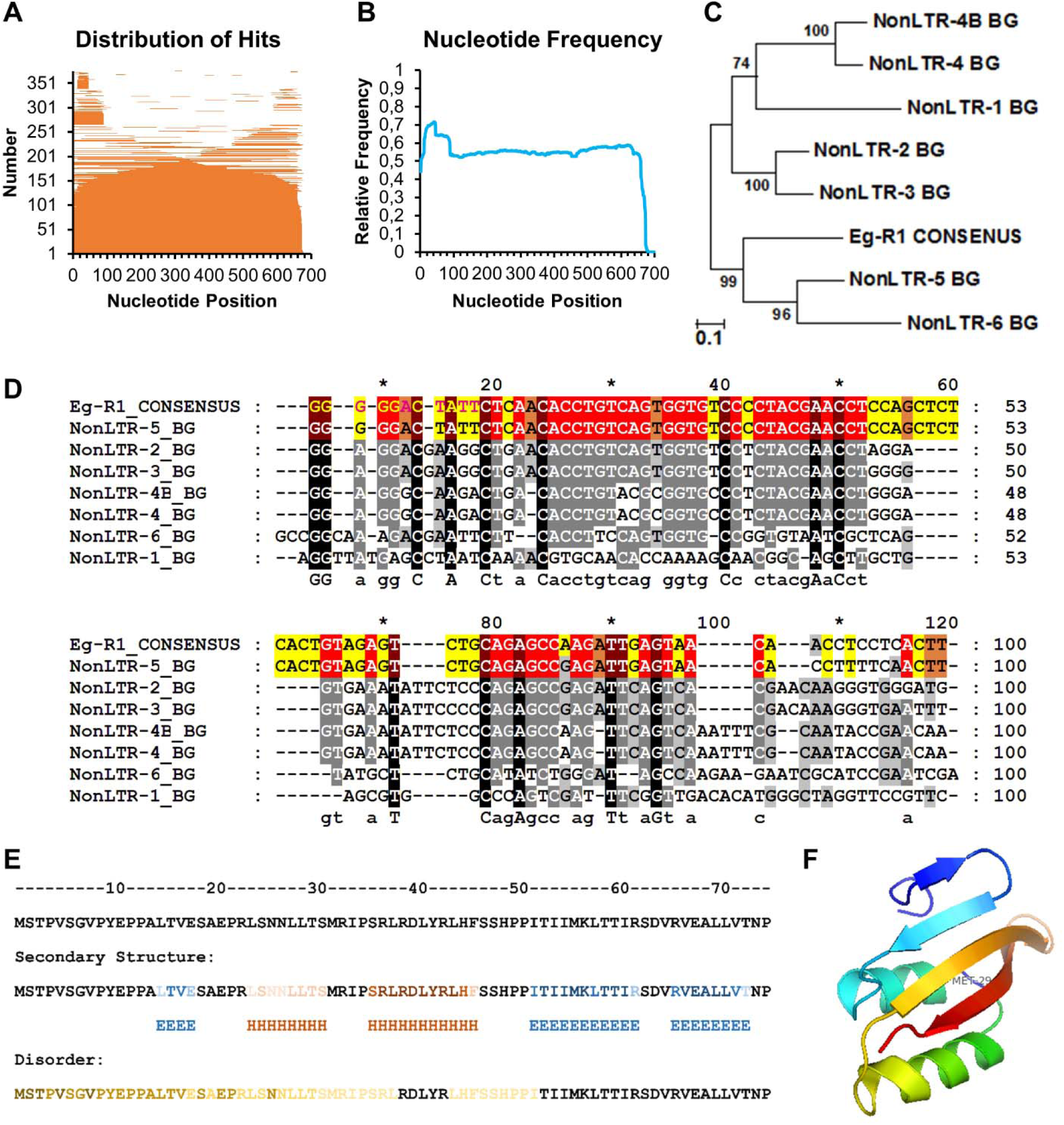
Genomic Insertion Size Distribution of Eg-R1, 5’-End Similarity of BG_Non-LTR Elements, Secondary and Tertiary Structure Predicton of ROPIP1. (A) Distribution of 376 BLASTn hits (BGH DH14 Genome v3b (contigs), blugen.org) using Eg-R1 (X86077.1) equipped with the 11 bp 5’ sequence extension as query. The hits are shown sorted in size and plotted to their respective position on the query. (B) The frequency of nucleotides matching to the query was calculated. No preferential insertion of any Eg-R1 part was observed. The increased nucleotide frequency within the first 88 nucleotides is likely due to the high sequence similarity of this sequence part with NonLTR-5_BG (see D). Together with the pattern in A this might support unequal crossing over mediated by Eg-R1 as suggested by [10]. (C) A 670 bp Eg-R1 consensus sequence was generated by nucleotide sequence alignment of 23 Eg-R1 full-length genomic insertions being surrounded by manually searched for target site duplications. Genomic Eg-R1 CONSENSUS contained the 11 bp 5’-sequence extension but lacked the poly (A) tail of the annotated Eg-R1 mRNA (X86077.1). 8 *Blumeria graminis* Non-LTR retrotransposons (BG_Non-LTRs) are deposited in Repbase Reports (2011, Volume11, Issue 9, [34]). EGRT1 Non-LTR Retrotransposon being identical to the annotated Eg-R1 sequence (X86077.1) was replaced by Eg-R1 CONSENSUS to calculate a phylogenetic tree by using the Maximum Likelihood method and 500 rounds of bootstrapping. (D) Nucleotide sequence alignment of the first 100 5’-nucleotides of the 8 BG_Non-LTRs. Eg-R1 CONSENSUS and NonLTR-5_BG shared a stretch of 88 identical nucleotides, except of one SNP, at their 5’-ends and were 93 % pairwise identical in the first 100 bp (identical residues are highlighted in color). All 8 BG_Non-LTRs were most identical (56 % pairwise identity) to each other within their first 100 bp. (E) ROPIP1 secondary structure prediction. Helix (H) and β-sheet (E) residues obtained from 4 algorithms provided by the Quick2D webserver [88] plus one obtained from the QUARK server [89] are shown combined. The darker the color the more algorithms predicted that residue. Results from protein disorder prediction using Quick2D [88] and DisEMBL [90] are depicted in a similar manner. Please note that the first Met of the ROPIP1 query is artificial. (F) *Ab initio* tertiary structure prediction of ROPIP1 obtained from the QUARK server [89]. The submitted query started with amino acid 5 (V) as depicted in E. Met 29 marks the start of ROPIP1-Cter in the model (amino acid position 32 in E). The estimated template modelling (TM-) score of the model was 0.3561 (TM-score ≥ 0.3: non-random structure).

**Fig. S5.**
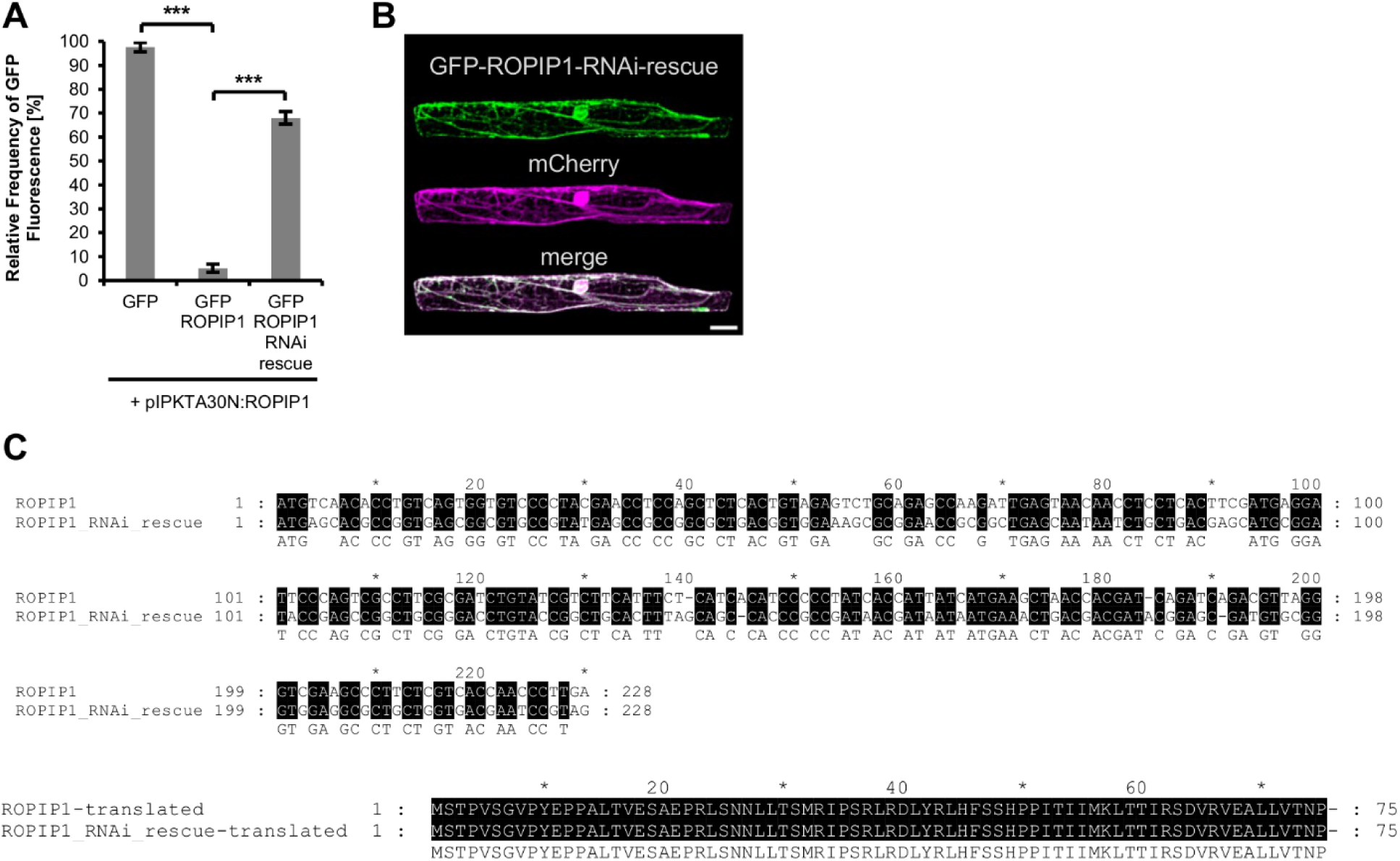
Test of Silencing Capacity of ROPIP1-RNAi and Sequence Alignment of ROPIP1 and ROPIP1-RNAi-Resuce. (A) Barley epidermal cells were transiently co-transformed with the RNAi vector pIPKTA30N-ROPIP1 (ROPIP1-RNAi) and pGY1-GFP or pGY1 -GFP-ROPIP1 or pGY1-GFP-ROPIP1-RNAi-rescue plus pGY1-mCherry as transformation marker. To test the silencing capability 1.0 μg/shot pIPKTA30N-ROPIP1 together with 1.0 μg/shot pGY1-GFP, or 1.0 μg/shot pGY1-GFP-ROPIP1 (see below), or 1.0 μg/shot pGY1-GFP-ROPIP1-RNAi-rescue and 0.5 μg/shot pGY1-mCherry each were co-transformed into barley epidermal leaf cells. MCherry expressing cells were categorized into GFP fluorescence visible or not at 36 hat by fluorescence microscopy. The relative frequencies of GFP-fluorescence were compared in a two-sided Student’s t-test. Transformed cells were identified by mCherry fluorescence and judged whether GFP fluorescence was visible or not at 36 hat by fluorescence microscopy. GFP-ROPIP1-RNAi-rescue was significantly less sensitive to ROPIP1-RNAi than GFP-ROPIP1. Bars are mean values of the relative frequencies of GFP to mCherry fluorescence from three independent experiments. In each experiment at least 150 cells per combination were scored. Error bars are ± SE. *** P ≤ 0.001 (Student’s t-test). (B) Confocal laser scanning microscopy whole cell projection of an exemplary barley epidermal cell showing translation of the GFP-ROPIP1-RNAi-rescue construct. The image was taken parallel to one repetition of (A). The co-transformed mCherry labeled the cytoplasm and the nucleoplasm. Co-localization is indicated by white pixels in the merge picture. Scale bar is 20 μm. (C) Nucleotide sequence and amino acid sequence alignment of the synthetic ROPIP1 RNAi rescue construct and ROPIP1. The codon usage of ROPIP1 was compared to the codon usage table of *Hordeum vulgare* subsp. *vulgare* in the codon usage database (kazusa.or.jp/codon/) using the graphical codon usage analyzer (gcua.schoedl.de/). Barley codons being most different, but not rare codons replaced the original codons of the ROPIP1 sequence. The identity of the two sequences was reduced to 64%. The wobble base exchanges of the ROPIP1 RNAi rescue construct were silent.

**Fig. S6.**
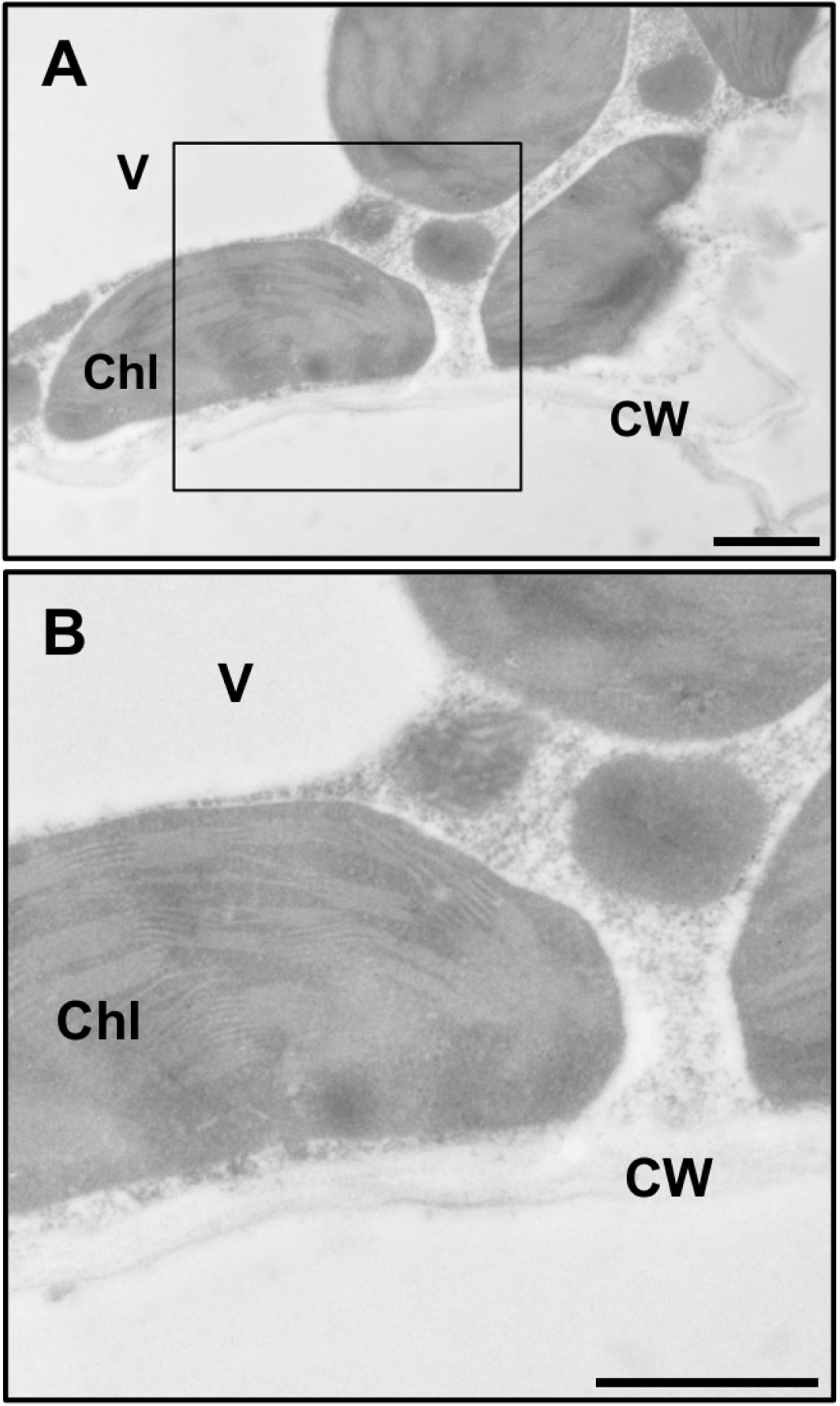
Immunogold Labeling of α-ROPIP1 in Mesophyll Cells of *Bgh*-Infected Barley Leaves. (A) Transmission electron micrograph of an ultrathin section of mesophyll cells of a Bgh-infected barley primary leaf at 3 dai. Gold-particles were almost absent throughout mesophyll cells after immunogold labeling of α-ROPIP1. (B) Detail picture of (A). Chl: chloroplast, CW: cell wall, V: vacuole. Scale bars are 1 μm.

**Fig. S7.**
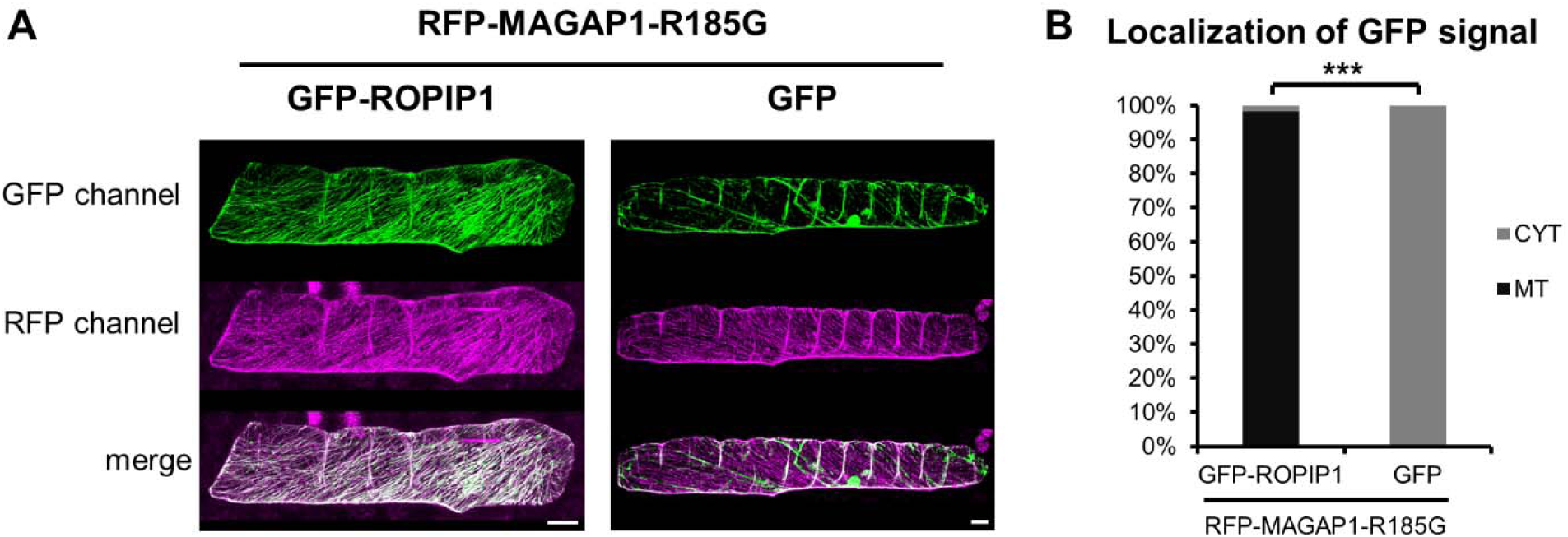
R185G Mutation of HvMAGAP1 Does Not Alter Microtubule Association of GFP-ROPIP1. Barley leaf epidermal cells were transiently transformed with RFP-HvMAGAP1-R185G plus GFP-ROPIP1 or GFP. HvMAGAP1-R185G lacks the catalytic arginine finger of its GAP domain. (A) Transformed cells were imaged as sequential whole cell scans by confocal laser scanning microscopy at 12 - 24 hat. (B) Maximum projections were categorized into GFP-fluorescence being located at microtubules (MT) or being absent from MTs but present in the cytoplasm (CYT). Bars represent relative frequencies of the categories derived from three independent replications. The respective absolute numbers per category of n = 59 GFP-ROPIP1 and n = 53 GFP expressing cells were compared in a X2 test. *** p ≤ 0.001 (X2). Scale bars are 20 μm.

**Fig. S8.**
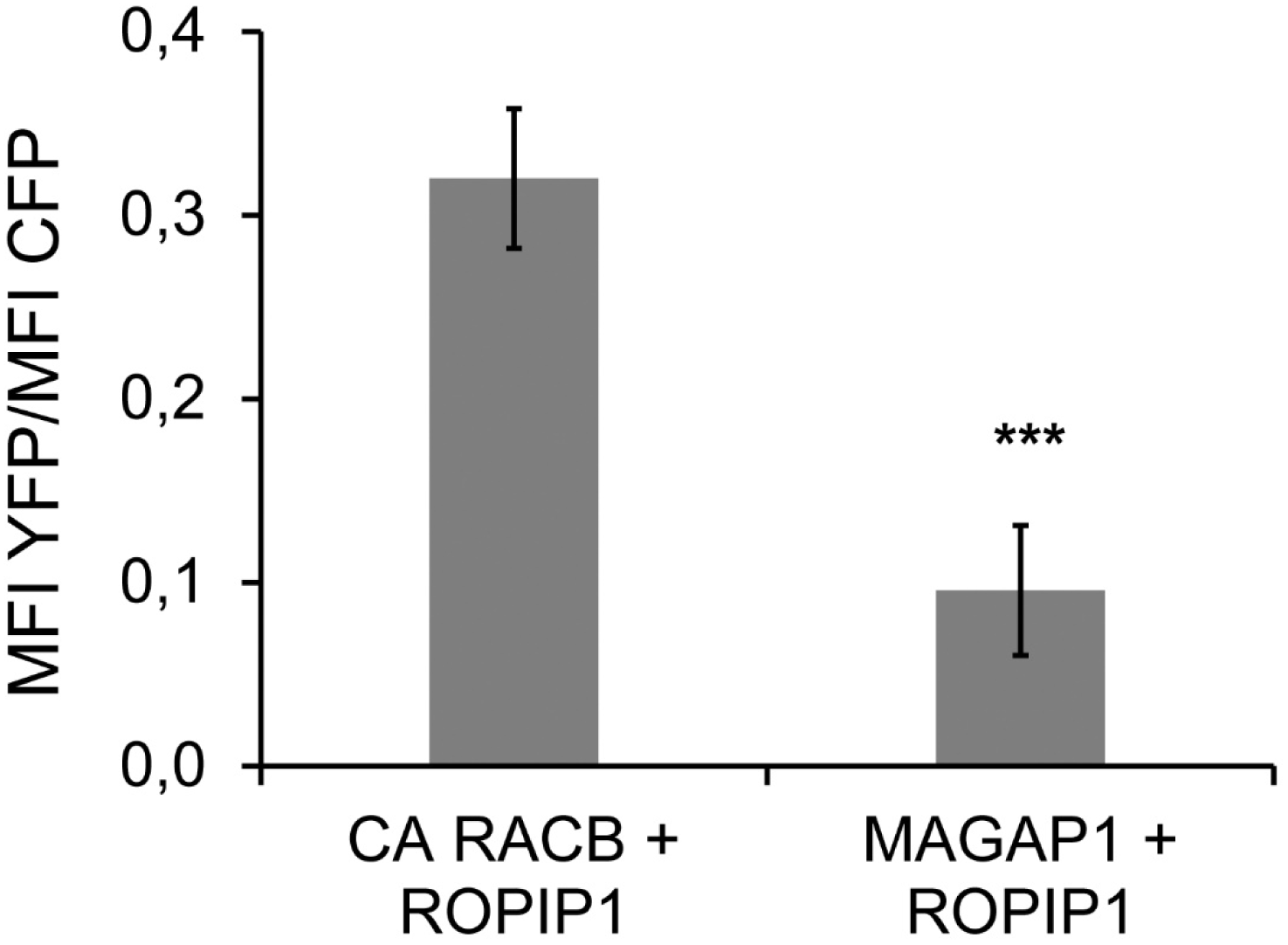
HvMAGAP1 Does Not Interact with ROPIP1 in a Split YFP Complementation Assay. (A) ROPIP1-YFP^N^ was transiently co-expressed with CA YFP^C^-HvRACB or YFP^C^-HvMAGAP1 and CFP as transformation marker in barley leaf epidermal cells. Ratiometric measurement of YFP fluorescence complementation as normalized to signals from co-expressed CFP. Error bars are ± S.E. Two-sided Student’s t-test (***; P ≤ 0.001).

**Table S1.**
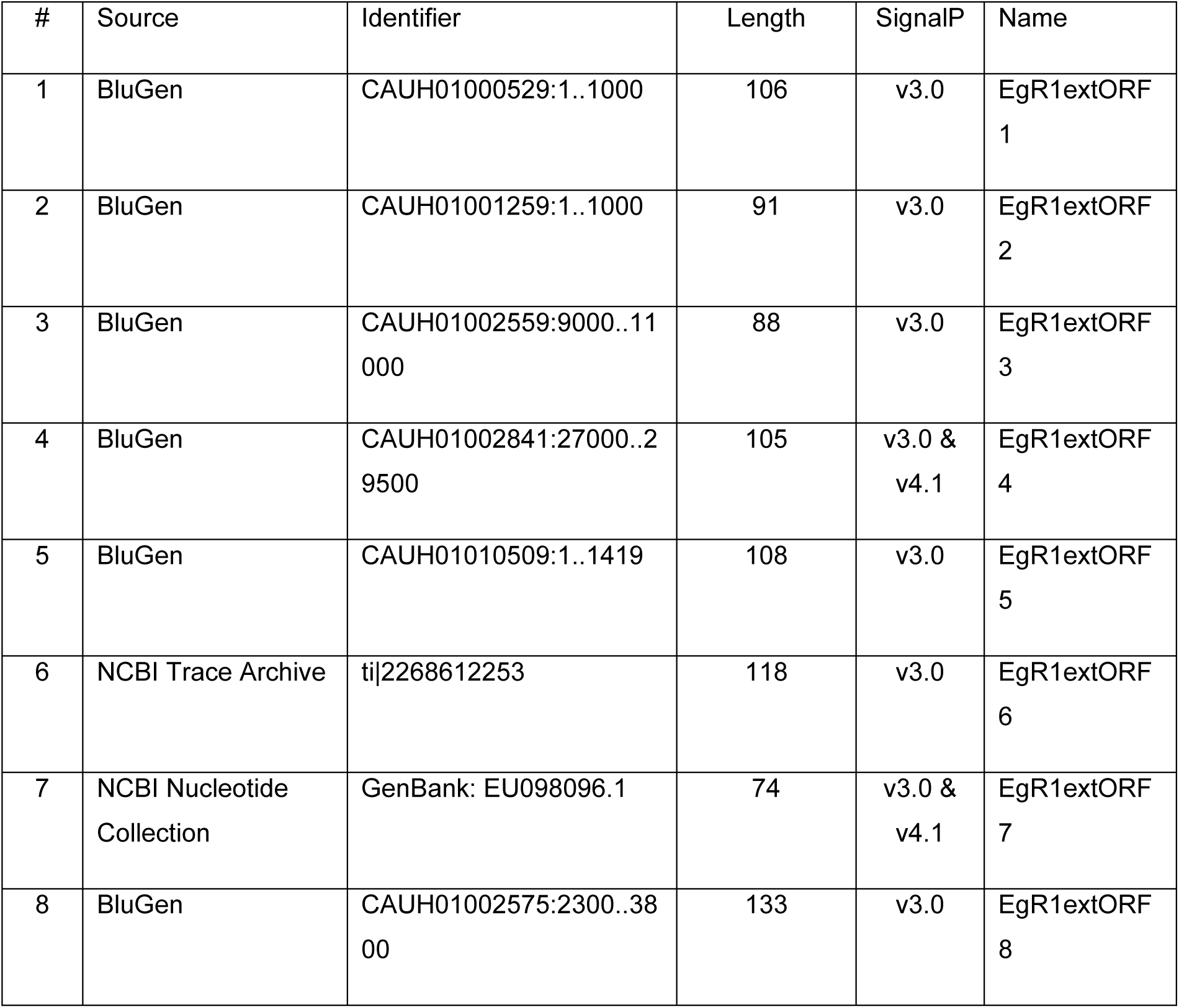
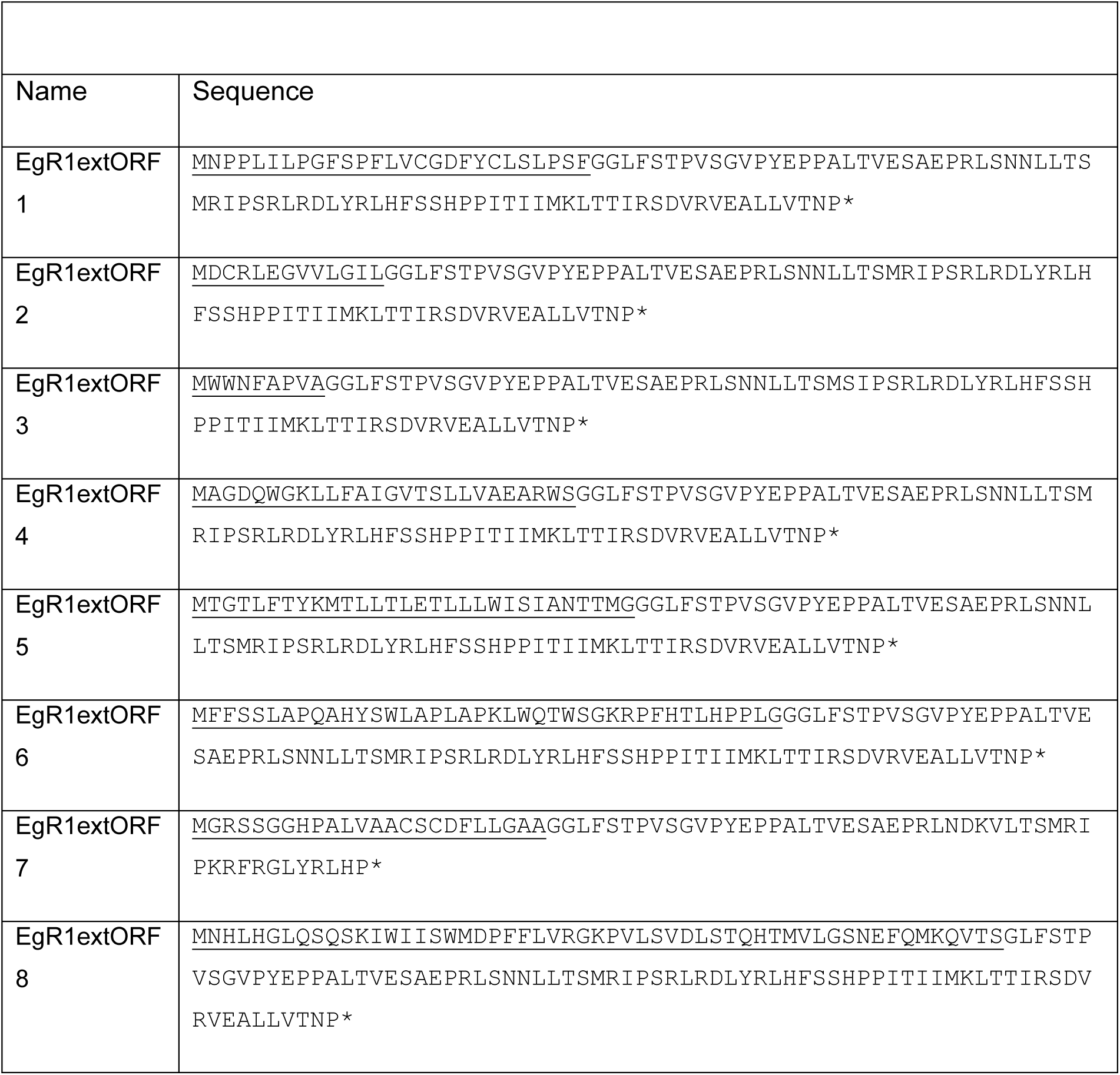
Genomic ROPIP1 Sequence Variants with Positive Signal Peptide Prediction. Signal Peptides were predicted using the SignalP server in the indicated version [86,87]. EgR1extORF1 till EgR1extORF5 were obtained from manually inspecting the topmost 45 BLASTn hits for 5’-elongated ROPIP1-ORFs using the ROPIP1 nucleotide sequence as query against the BGH DH14 Genome v3b database of *Bgh* genomic DNA contigs (blugen.org). EgR1extORF6 was found among the manually inspected 10 topmost BLASTn hits of the ROPIP1 query against the NCBI Trace archive database ‘Blumeria graminis f sp hordei WGS’ that contains *Bgh* whole genome shotgun raw reads. The 3’-truncated EgR1extORF7 was found in vicinity to an AVRa10-like effector protein by using the ROPIP1 nucleotide query against the NCBI non-redundant nucleotide collection database. EgR1extORF8 contains a putative intron between the predicted signal peptide and the ROPIP1 sequence. Sequence extensions of ROPIP1 contributing to a positive signal peptide prediction are underlined. EgR1extORF2, EgR1extORF3, EgR1extORF5 and EgR1extORF8 were amplifiable from genomic DNA prepared from *Bgh* race A6-infected barley leaves using gene-specific primers.

**Table S2:**
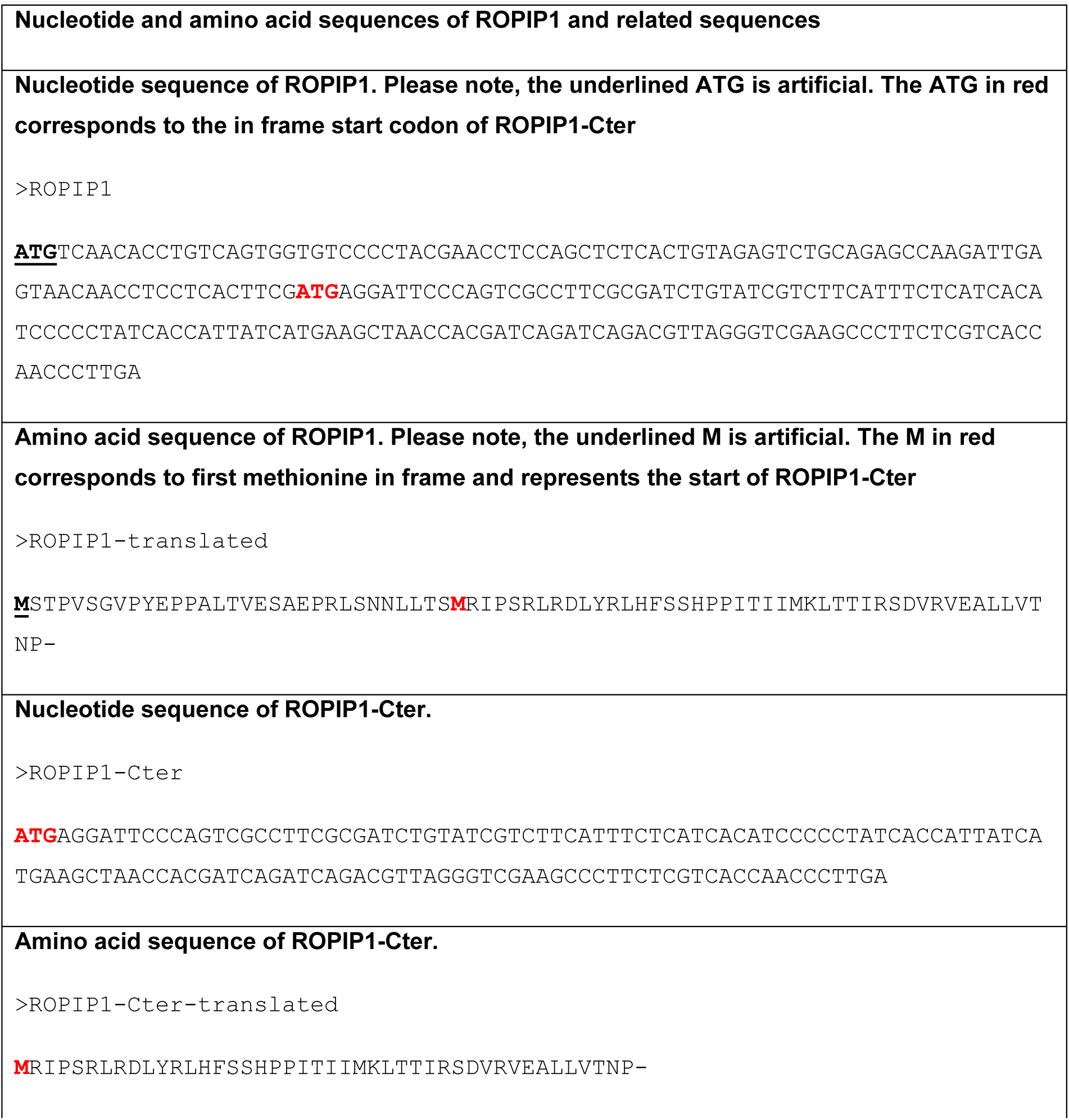
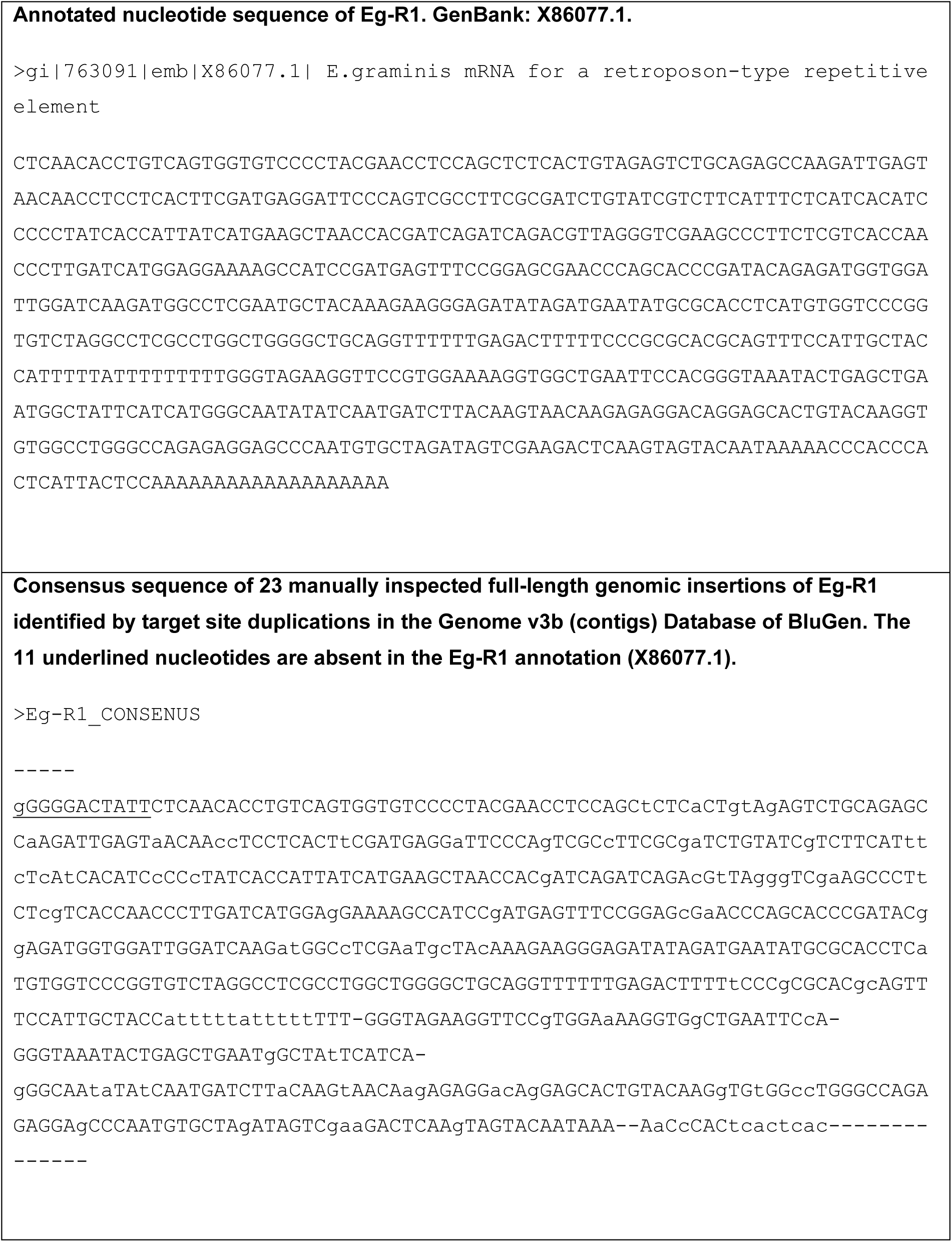
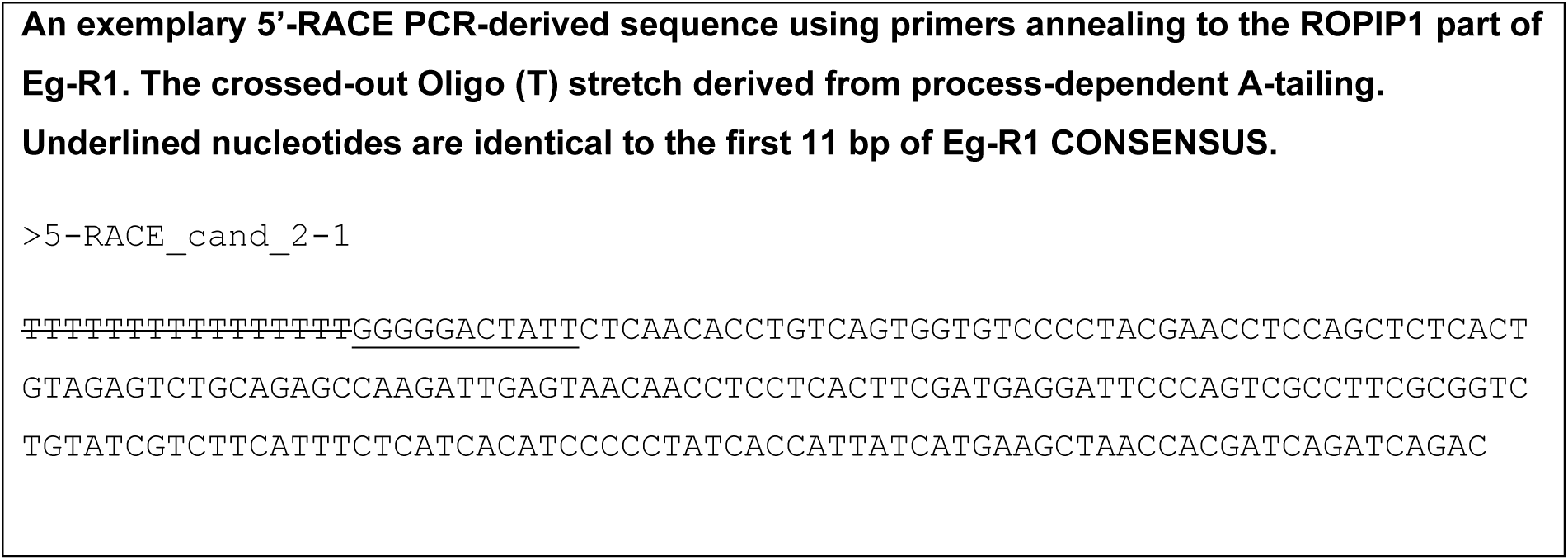
Nucleotide and Amino Acid Sequences of ROPIP1 and Eg-R1.

**Table S3.**
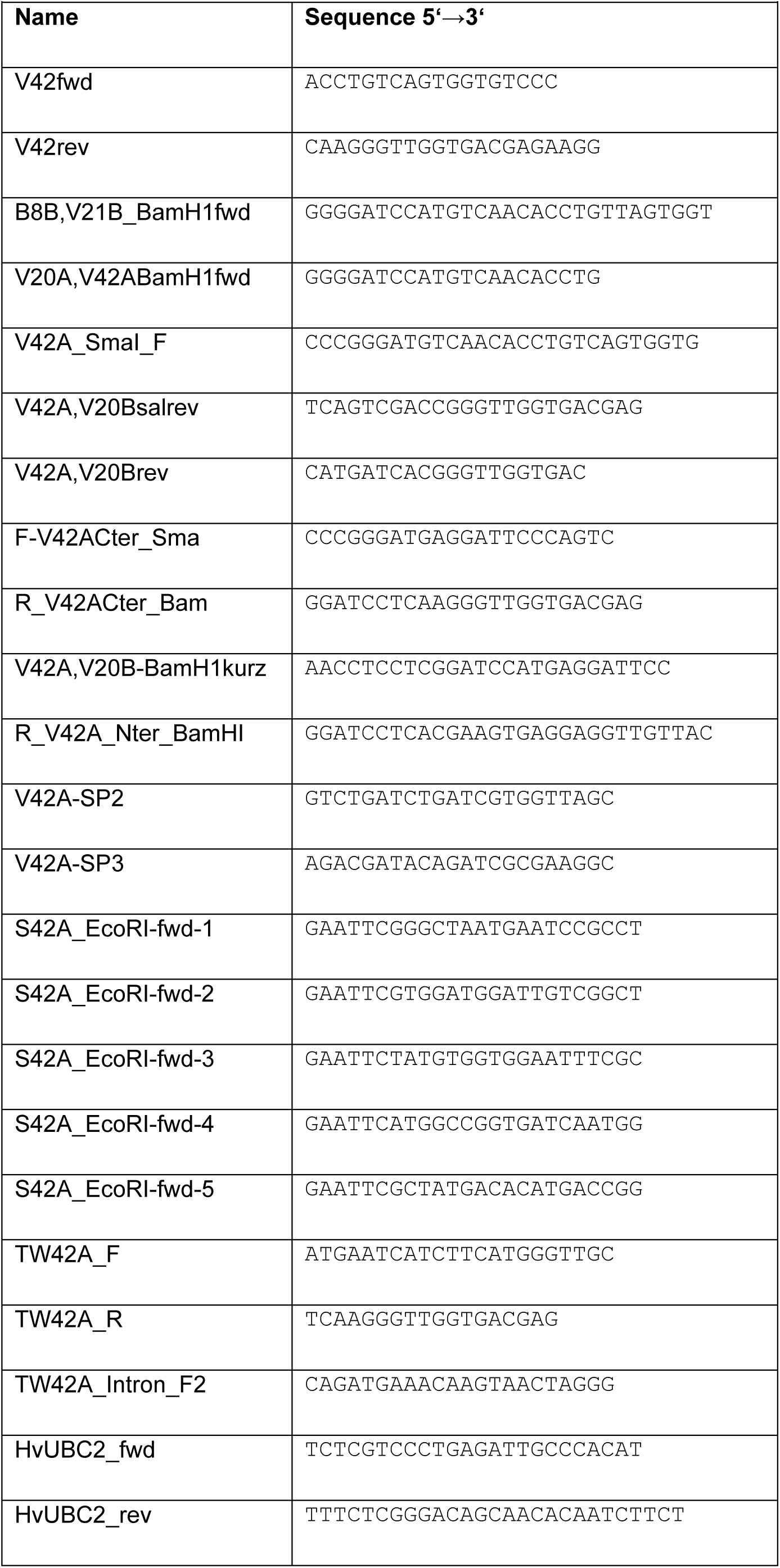
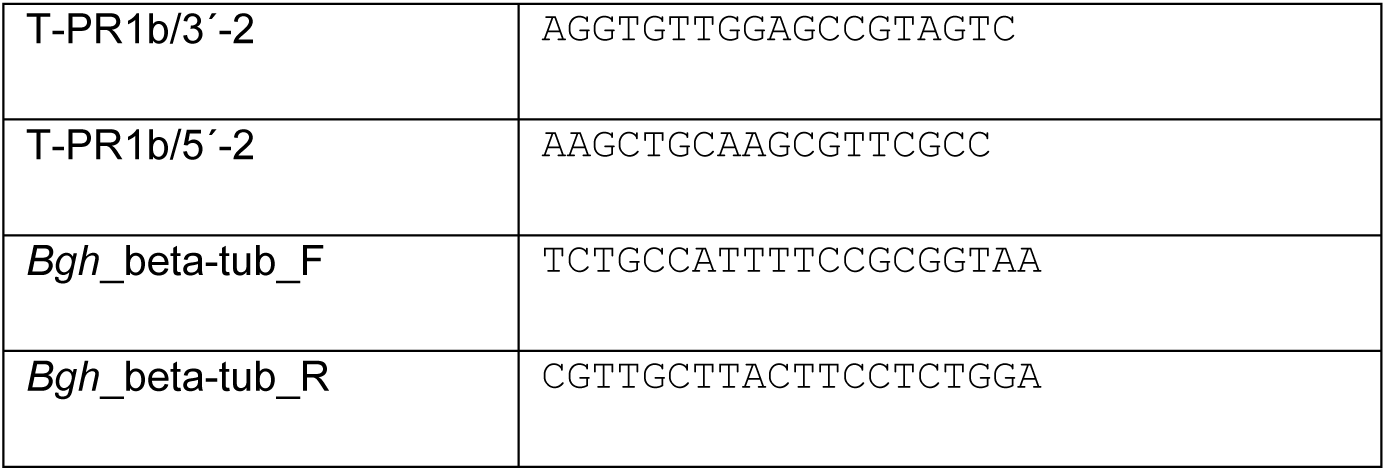
List of Oligonucleotides Used in this Study.

